# Multifaceted analysis of training and testing convolutional neural networks for protein secondary structure prediction

**DOI:** 10.1101/2020.01.17.911065

**Authors:** Maxim Shapovalov, Roland L. Dunbrack, Slobodan Vucetic

## Abstract

Protein secondary structure prediction remains a vital topic with improving accuracy and broad applications. By using deep learning algorithms, prediction methods not relying on structure templates were recently reported to reach as high as 87% accuracy on 3 labels (helix, sheet or coil). Due to lack of a widely accepted standard in secondary structure predictor development and evaluation, a fair comparison of predictors is challenging. A detailed examination of factors that contribute to higher accuracy is also lacking. In this paper, we present: (1) a new test set, *Test2018*, consisting of proteins from structures released in 2018 with less than 25% similar to any protein published before 2018; (2) a 4-layer convolutional neural network, *SecNet*, with an input window of ±14 amino acids which was trained on proteins less than 25% identical to proteins in *Test2018* and the commonly used *CB513* test set; (3) a detailed ablation study where we reverse one algorithmic choice at a time in *SecNet* and evaluate the effect on the prediction accuracy; (4) new 4- and 5-label prediction alphabets that may be more practical for tertiary structure prediction methods. The 3-label accuracy of the leading predictors on both *Test2018* and *CB513* is 81-82%, while *SecNet*’s accuracy is 84% for both sets. The ablation study of different factors (evolutionary information, neural network architecture, and training hyper-parameters) suggests the best accuracy results are achieved with good choices for each of them while the neural network architecture is not as critical as long as it is not too simple. Protocols for generating and using unbiased test, validation, and training sets are provided. Our data sets, including input features and assigned labels, and *SecNet* software including third-party dependencies and databases, are downloadable from dunbrack.fccc.edu/ss and github.com/sh-maxim/ss.

## Introduction

The prediction of secondary structure—alpha helices, beta sheet strands, and coil regions— is a long-standing problem in computational biology [1]. Linguistically, the secondary structure prediction problem is a translation of a word formed from a 20-letter amino-acid alphabet to a word of the same length that designates the secondary structure. The commonly used *DSSP* program [2] designates the secondary structure of experimental structures with an 8-letter alphabet (*H*, *B*, *E*, *G*, *I*, *T*, *S*), which can be reduced to a smaller alphabet (commonly *H*, *E*, *C*) according to defined rules. The results of secondary structure prediction have often been used as inputs to tertiary structure prediction methods [3-9], estimation of folding rates [10], prediction of solvent exposure of amino-acid residues [11-13], prediction of beta-turn locations and types [14-16], discrimination of intrinsically disordered regions [17, 18], accurate multiple sequence alignment [19-22], protein function prediction [23, 24], and prediction of missense mutation phenotypes [25].

The problem of predicting secondary structure from sequence has a long history, starting in 1965 [26]. Over a period of more than 50 years, the accuracy has gradually increased due to an increase in the number of experimentally determined structures in the Protein Data Bank (PDB), the amount of sequence information available for the determination of features derived from multiple sequence alignments, computational power, and the evolution of machine learning algorithms. Structures in the PDB provide information for deriving the ground-truth secondary structure labels used for training and testing of predictive models. It currently contains more than 150 thousand experimental structures and about 15 thousand non-redundant proteins (at 25% sequence identity) [27]. The number of available protein sequences in sequence databases has dramatically increased from 5 million in 1999 when the widely-used *PSIPRED* was published [28] to 168 million today.

Secondary structure prediction methods and our ability to benchmark them have been evolving and improving too. Several reviews [1, 19, 29-33] provide an excellent source of the history of improvements in prediction methods. These methods have been divided into a set of four generations of methods covering the 50-year time period from 1960 to 2010 [1, 30, 34]; we also suggest a new fifth generation from the 2010s until today and describe it in more detail.

The *first generation* of methods—such as *C+F* [35], *Lim* [36], and *GORI* [37]—from the 1960s and 70s relied on single amino-acid preferences for the *H*, *E*, and *C* secondary structure types. In 1983 the original accuracy of 65-70% reported by the first-generation methods was revised downwards to 48-56% [38]. Dickerson et al. were the first to demonstrate that evolutionary information was helpful for secondary structure prediction [39]. However the evolutionary approach during this period was limited by the small number of homologous protein sequences available for any target.

The *second-generation* methods from the 1980s and early 1990s—such as *Schneider* [40], *ALB* [41], *GORIII* [42], *COMBINE* [43], and *S83* [44]—utilized statistical analysis of a stretch of adjacent residues to predict secondary structure of a central residue [30, 34]. These were based on statistical information, sequence patterns, physiochemical properties, neural networks (NN), graph theory, multivariate statistics, expert rules, and nearest-neighbor algorithms [34]. The second-generation methods benefited from larger sequence databases and usage of evolutionary information. For instance, Zvelebil et al. incorporated evolutionary information in their method by predicting secondary structure for each protein in an alignment and then reporting an average prediction [45]. However, the second generation methods saturated at a low 3-label accuracy of 58-63% and experienced two problems: 1) beta strands were predicted at very low accuracy levels of 28-48%, marginally better than random; 2) predicted helices and strands were too short to be practical [34].

The methods of the *third generation* from the mid 1990s—such as *PHD* [46], *LPAG* [47], *NSSP* [48]—achieved a 10% improvement in the 3-label accuracy over second-generation methods, leading to 68-72% accuracy. Methods from the late 1990s to the early 2000s, including *PSIPRED* [49], *JPred2* [50], *SSpro* [51], and *PROF* [52], further reached 75-77%. The success came from three factors [30, 34]: 1) use of evolutionary information encoded in a multiple-sequence alignment (MSA) profile or position specific scoring matrix (PSSM) as direct inputs to a prediction program; 2) larger databases than for the second-generation methods; and 3) more advanced algorithms. The algorithms based on neural networks [30, 46, 49, 53, 54] demonstrated the greatest improvement.

Inspired with evolutionary information features as input in past applications, the *fourth-generation* methods from mid-late 2000s further utilized a variety of additional input features [33] including penta-peptide statistics [55, 56], conserved domain profiles [57], frequent amino-acid patterns [58], predicted torsion angles [59, 60], predicted residue contact maps [61], predicted residue solvent accessibility [62], predicted tertiary structure [62, 63], and predicted pseudo-energy parameters for helix formation [64]. Most of these individually led to small improvements. For example, Meiler et al. suggested seven representative amino-acid properties as input and showed their tenfold cross-validated accuracy increased only by 0.5% from 77% [65]. The predicted residue solvent accessibility improved the 3-label accuracy by 3% [66]. The predicted dihedral angles helped by 2% [67]. However, these features together did not improve the 3-label accuracy beyond 80% [68]. Some of the fourth-generation methods—such as *HYPROSP* [69], *PROTEUS* [70], *MUpred* [71], and *DISTILL* [72]—took advantage of structural fragments or templates of homologous sequences to attempt a breakthrough. It only brought a modest 3% improvement and did not raise the 3-label accuracy beyond 80% with or without the additional input features.

With greatly increased computing capabilities (cost, memory, CPU, and GPU) and exploding popularity of deep neural networks, the *fifth-generation* methods from 2010 up to the present: 1) revisited the template-based approach [73, 74]; 2) included sequence alignment parameters from hidden Markov models (HMM) as input features [75-78]; 3) were designed with more advanced neural network architectures (convolutional neural networks (CNN), bidirectional recurrent NNs such as long short-term memory, and residual NNs and their combinations) [74- 82]; 4) more painstakingly employed an ensemble of models with an average vote [74, 76, 78, 81, 83, 84]; and 5) in some cases, trained on predicted residue contacts, residue surface accessibility, Cα-atom exposure, backbone torsion angles, and 3-label secondary structure predictions from other programs [84]. The template-free [73, 75-83] methods reported accuracies from 82 to 87%, a substantial improvement over the fourth-generation accuracy of 80%. The theoretical 3-label accuracy limit is 90-95% [1, 30, 72, 84-86], based on the rate at which different programs such as *DSSP* and *Stride* [87] assign the same secondary structure to experimental coordinates.

In 2015, the fifth-generation method *SPIDER2* [82, 88] utilized three iteratively connected CNNs, each consisting of three hidden layers, achieving a 3-label accuracy of 82%. *Jpred4* [76] had the same 82% accuracy in the same year by combining several CNNs trained on the same multiple sequence alignments presented to the networks in different ways. One year later in 2016, *DeepCNF* [79] achieved a 2% accuracy improvement to 84% by combining a 5-7 layer CNN with conditional random field as an additional layer. The CNN was constructed in a funnel style by enforcing the same weights in neighboring input and hidden nodes and thus limiting the total number of parameters to train and allowing for longer-range sequence information. The final conditional random field layer was used to account for correlation of adjacent residues. In 2017, the authors of *SPIDER2* upgraded their previous method to a new NN architecture in *SPIDER3* [75], relying on four hidden layers with the first two bidirectional recurrent NN (BRNN) layers followed by two fully connected layers, and reporting an 84% 3-label accuracy, an improvement of 2% over *SPIDER2*. Also in 2017, *FSVM* [73] was reported with an 83% template-free accuracy achieved with a fuzzy support vector machine. In 2018, *MUFOLD-SS* [77] reported a 3-label accuracy of 84% with a neural network relying on nested inception models which are nested networks of several parallel CNNs that proved to be state-of-the-art in image recognition. At the same time, *PORTER5* [78] achieved the same 84% accuracy by employing an ensemble of BRNNs. In 2018, another method, *PSRSM* [83] reported an improvement of 1-2% to 85-86% accuracy; however, in an independent study the reported accuracy was revised downward to 82% [84]. The *PSRSM* publication does not mention a sequence similarity exclusion between training and test sets, which could be a cause for this overestimation. The 2018 method, *CNNH_PSS* (a 5- layer multiscale CNN with highways between neighboring layers), was only trained for 8 labels and reported 70% accuracy [80]. Another 2018 method, *eCRRNN*, an ensemble of 10 networks based on a combination of convolutional, residual, and bidirectional recurrent NNs), claimed the highest 8-label and 3-label accuracies to-date of 74% and 87% respectively [81].

The fifth-generation methods [73-84] all depend on multiple sequence alignments of the target sequences, while several of these methods also rely on templates [73, 74], i.e., real protein structures from a training set consisting of experimental coordinates of protein fragments. These “template-based” methods can achieve higher accuracies, in the range 86-93%, than template-free methods. However, homologous structures are not available for many proteins that would be targets of secondary structure prediction, and such methods are not solutions to the general secondary structure prediction problem. The accuracy of the template-based methods drops from 86-93% to 80-83% without templates and further down to 74-77% without homologous sequences [73, 74, 84]. Similarly, the accuracies of the template-free MSA-based methods drop from 82-84% to 73-75% without the use of homologous sequences [79].

The template-based *SSpro5* from 2014 [74] utilizes a *BLAST* [89] search of the PDB to find similar sequence fragments of at least length 10 to a target sequence and reports the most common *DSSP*-assigned class in the set of proteins selected for a given position. When no similar sequences or no dominant *DSSP* class in the similar sequences are found, secondary structure prediction is based on an ensemble of 100 BRNNs trained on the data set. With templates, *SSpro5* achieves 93% 3-label accuracy; without templates *SSpro5* has only 79-80% accuracy. *FSVM* [73] described above can also run in a template-based mode by using the same sequence-based structural similarity concept as in *SSpro5*. In this mode, *FSVM*’s 3-label accuracy increases from 83% to 93%.

One common issue that has recurred repeatedly during the history of secondary structure prediction is that reported accuracies have not always been upheld when methods were applied to new benchmark test sets [30, 34, 38, 46, 84, 90]. There are a number of reasons for this. The quality and size of test sets for benchmarking have sometimes proven inadequate and often the test sets are not distinct enough from the training sets so the methods are over-trained [34, 46, 90, 91]. In some cases, training was performed on sequence sets representing the entire sequence space of the PDB leading to very similar proteins in training and testing sets [83], or a test set was inadvertently used multiple times for development decisions. It is challenging to compare programs on common benchmark sets because the programs are trained on different sets of structures, some of which may be in any particular benchmark. For example, the *CB513* data set [91] has been used repeatedly [73, 77, 79-81], even though related proteins may be used in the training data for a new program [74, 75, 78, 84].

In this paper, we investigate factors that contribute to the accuracy of template-free protein secondary structure with (1) an ensemble of 10 simple traditional 4-layer CNNs and (2) rigorously defined and independent training, validation, and test sets. Our test set, *Test2018,* consists of proteins whose structures were determined in 2018 that do not share more than 25% sequence identity with any structure of any resolution or experiment type deposited before January 1, 2018. This enables us to compare our program, *SecNet*, with methods that were trained prior to the beginning of 2018. While there has been only ∼5% improvement from third-generation methods of the early 2000s, we demonstrate an accuracy increase of 2-3% in both the 3- and 8-label accuracies compared to two recently developed deep learning models which had reported the highest accuracy on the popular *CB513* data set available since 1999. We demonstrate that one of these methods, *eCRRNN*, with the best reported 8-label accuracy of 74% and 3-label accuracy of 87% to date, achieves accuracies of 70.8% and 81.8% respectively for the 8 and 3 labels on our *Test2018* test set while our program achieves 73.0% and 84.0% respectively. We describe how to benchmark existing or future methods against the new test set (or similarly constructed test sets in the future) and what mistakes to avoid during training and testing.

Through an ablation study that followed *SecNet* development, we investigated factors that are important for high-accuracy prediction such as method complexity, types of input features, window size, database source and size, alignment parameters, and training hyper-parameters. For example, our results show that CNNs do not benefit in terms of overall accuracy beyond 15 residues away from a prediction label. We discuss the prediction practicality of secondary structure labels for protein tertiary structure prediction, and propose new 4 and 5 prediction label schemes that should be more useful for structural biology.

## Results

### Data sets for training, validation, and testing secondary structure prediction methods

We produced separate sets of structures from the PDB for our training, validation, and test data sets with the protocol shown in Fig 1 and fully described in *Methods*. Our aim was to make the protocol reproducible, so that it can be reused for creation of new test sets in the future. Because we wanted to compare our secondary structure prediction program with the results in earlier publications, we developed a test set that would not have proteins in the training and testing data of earlier programs. Similar approaches for deriving test sets have been used by others [84].

**Fig 1.**
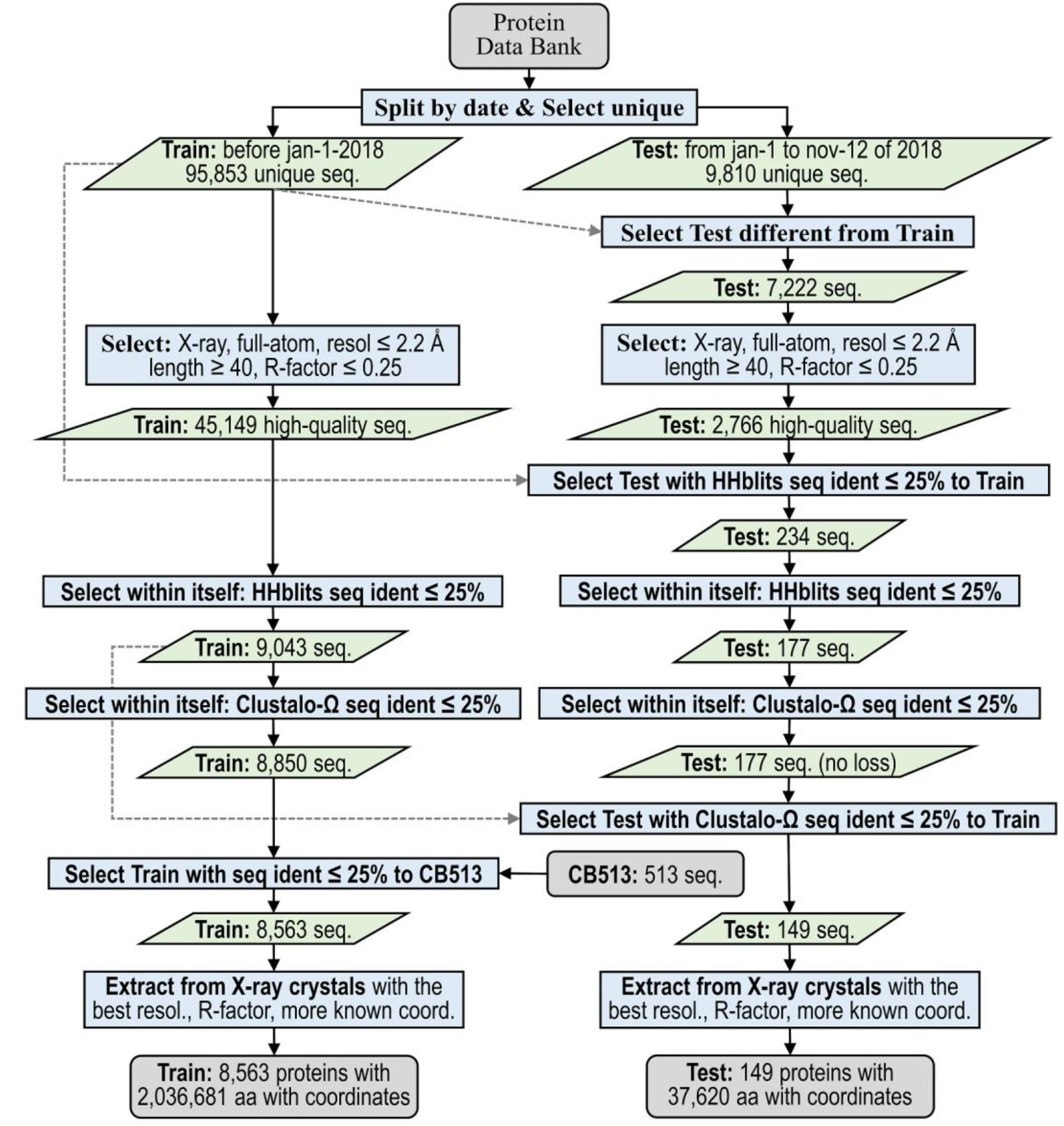
Protocol flowchart for preparation of 2018 unbiased test and training sets (*Set2018* dataset). Protocol input and output are shown with gray rounded rectangles. All action processes are in blue rectangles. Green parallelograms represent intermediate input and output data. The split date for *Set2018* training and test sets is Jan 1, 2018. By using two sequence alignment programs, the protocol removes from the test set any proteins with more than 25% sequence identity to any previously published PDB structure of any experimental type, resolution, or quality. The test set guarantees unbiased accuracy estimation for our prediction method, *SecNet*, and any previous software trained and validated on proteins released before Jan 1, 2018.

To develop the test set, we placed all 9,810 unique protein chains in PDB entries of any resolution or experimental type released between January 1 and November 12, 2018 into a starting set. We filtered the test proteins to satisfy a 2.2 Å resolution threshold, a 0.25 R-factor cutoff consistent with this resolution [92], and a minimal chain length of 40 residues. From this reduced set, we additionally removed any sequence with more than 25% sequence identity to any of the 95,853 unique sequences in structures released before January 1, 2018 of any resolution, any experiment type, and any quality-control characteristics. This similarity exclusion took place if either of two programs, *HHblits* [93] or *Clustal-Omega* [94], calculated a pairwise sequence similarity above the 25% cutoff. Finally, we applied the same pairwise sequence similarity cutoff of 25% according to the same two alignment programs within the test set itself. In the last step, when several similar proteins had to be removed, the algorithm kept proteins with a higher resolution or better R-factor or more residues with known coordinates in that order [27].

For the training/validation set, the same resolution, R-factor, length, and sequence identity criteria were applied to the sequences and structures that were released by the PDB prior to 2018. Chains with sequence identity ≥ 25% with any protein in the popular *CB513* data set were also removed from the set so that *CB513* could also be used in testing (see below). The resulting set was split with a 9:1 ratio to produce the training and validation sets. We applied *DSSP* to the chains in each data set, producing training, validation, and test sets respectively that have 7707, 856, and 149 protein chains and 1830, 207, and 38 thousand amino acids of labeled secondary structure (Table 1 and Table A in S1 Supporting information). We refer to the three data sets collectively as *Set2018*, and the testing set individually as *Test2018*. We provide *Set2018* (S2 Data set) including the PDB entry and chain, the full-length original and unmodified sequences using a standard 21- letter notation (with modified amino acids represented with single letters of their unmodified counterparts where possible; other non-standard amino acids are represented by “X”), and the *DSSP* codes (including the symbol “X” for residues with missing backbone coordinates).

**Table 1.** Statistics for the Set2018 training, validation, and testing data sets.

| Set2018 |  |  | Proteins | Perc. of amino acids | No of amino acids |
| --- | --- | --- | --- | --- | --- |
| All = Train + Valid + Test |  |  | 8,712 | - | - |
| whole sequence |  |  | - | 100.0 | 2,217,707 |
| with coordinates |  |  | - | 93.5 | 2,074,301 |
| w/o coordinates |  |  | - | 6.5 | 143,406 |
| All with coordinates |  |  | 8,712 | 100.0 | 2,074,301 |
| H | α-helix | | - | 34.1 | 707,050 |
| E | strand |  | - | 22.3 | 463,310 |
| C | coil |  | - | 19.4 | 401,922 |
| T | turn |  | - | 11.1 | 229,566 |
| S | bend |  | - | 8.2 | 169,326 |
| G | 3<sub>10</sub> helix or helix-3 |  | - | 3.9 | 80,536 |
| B | β-bridge | | - | 1.1 | 22,245 |
| I | π-helix or helix-5 | | - | 0.017 | 346 |
| Training set |  |  | 90% split of (All – Test) by proteins |  |  |
| Any |  |  | 7,707 | 100.0 | 1,830,092 |
| C | coil | Rule #1 | - | 38.6 | 706,364 |
| H | helix |  | - | 38.0 | 695,750 |
| E | sheet |  | - | 23.4 | 427,978 |
| Validation set |  |  | 10% split of (All – Test) by proteins |  |  |
| Any |  |  | 856 | 100.0 | 206,589 |
| C | coil | Rule #1 | - | 38.7 | 79,985 |
| H | helix |  | - | 37.9 | 78,261 |
| E | sheet |  | - | 23.4 | 48,343 |
| Test set |  |  | Fixed, 2% of All |  |  |
| Any |  |  | 149 | 100.0 | 37,620 |
| C | coil | Rule #1 | - | 38.5 | 14,465 |
| H | helix |  | - | 37.0 | 13,921 |
| E | sheet |  | - | 24.5 | 9,234 |

#### SecNet: a 4-layer convolutional neural network for protein secondary structure prediction

We describe a traditional CNN with only 4 hidden layers (Fig 2), trained on our pre-2018 training set from *Set2018*. The validation set was used multiple times to try different neural network architectures, to select features having a positive impact on prediction, and to optimize feature and training parameters. After the best model based on the highest validation accuracy during training was saved, the program was only once benchmarked on our unbiased *Test2018* test set. The 92 input features of our template-free method consist of one-hot encoding of an amino-acid sequence, two *PSI-BLAST* profiles (after 1 and 3 rounds of search on Uniprot90), and HMM parameters derived from a multiple sequence alignment of a target sequence and hits from the Uniprot20 sequence database [93]. We underline that our predictive model is a simple, traditional CNN, which is in contrast to the recent trend of using more and more complex methods and NN architectures [74-82]. It employs an input window of only 29 amino acids. The number of input parameters from the 2018 training set is 1,830,092 amino acids x 92 features per amino acid = 168,368,464. Our NN has 2,397,960 of trained parameters with a ratio of input to trained parameters equal to 70.2 = 168,368,464 / 2,397,960. A detailed description of our CNN and input features is provided in *Methods*.

**Fig 2.**
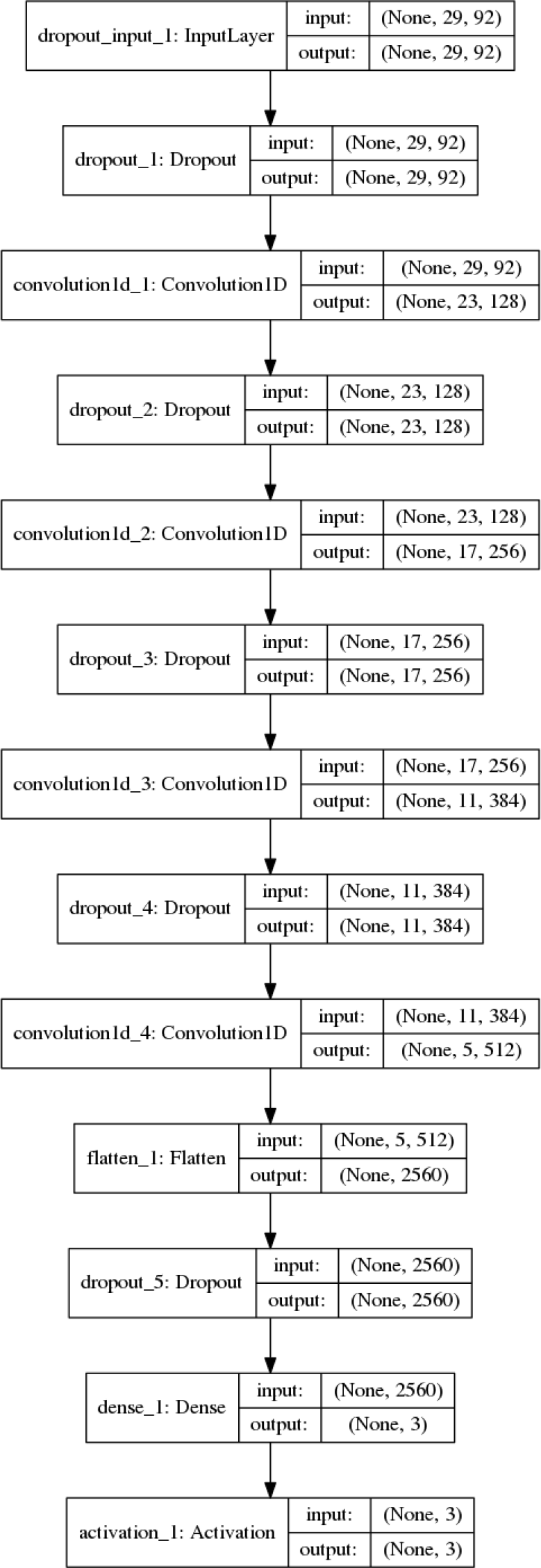
Architecture of *SecNet*. *SecNet* is a traditional CNN with an input layer, 4 hidden convolutional layers, and a dense layer with dropout regularization layers in between. The input layer reads a 29 x 92 matrix representing a sequence window of 29 amino acids centered on the one subject to prediction and 92 input features for all 29 positions. 92 features = one-hot encoding of 22 amino-acid types + 2 rounds x 20 *PsiBlast* profile values + 30 HMM alignment parameters. Each box encloses a linear dimensionality and number of features (2^nd^ and 3^rd^ values in parentheses) in input and output of each layer. The total number of parameters to train is about 2 million. The activation layer returns 3 probabilities for the 3 labels: *H*, *E*, and *C* of the central amino acid which are calculated with a *softmax* activation function. The label with the highest estimated probability is the predicted label.

We first tested the accuracy of predicting all 8 labels from *DSSP* (*H, E, C, G, I, S, T, B*); the 8-label accuracy is 73.0% on *Test2018*. To develop a program that is trained and tested on three labels (*H, E*, *C*), we need a rule to convert the 8-letter alphabet to the 3 letters. The most common rule, which we refer to as *Rule #1*, is (*H*, *G*, *I*) → *H*, (*E*, *B*) → *E*, (*T,S,C*) → *C* [95]. A less common rule, which we refer to as *Rule #2*, has a simpler definition: (*H*) → *H*, (*E*) → *E*, (*G,I,B,T,S,C*) → *C* has been used in some programs [53, 76, 81, 96-99]. Several published methods did not clearly define which 3-label rule was used [73, 77, 78, 82]. If we use *Rule #1* on the training, validation, and testing data, we achieve a 3-label accuracy of 84.0% on *Test2018*. The confusion matrices, true positive rates (recalls), positive predictive values (precisions), false negative rates, and false discovery rates for the 8-label and 3-label Rule #1 predictions are presented in Tables 2 and 3 respectively. When *SecNet* is trained, validated, and tested on data derived using *Rule #2*, which is easier to predict [28, 91], it has a higher accuracy of 86.0%.

**Table 2.**
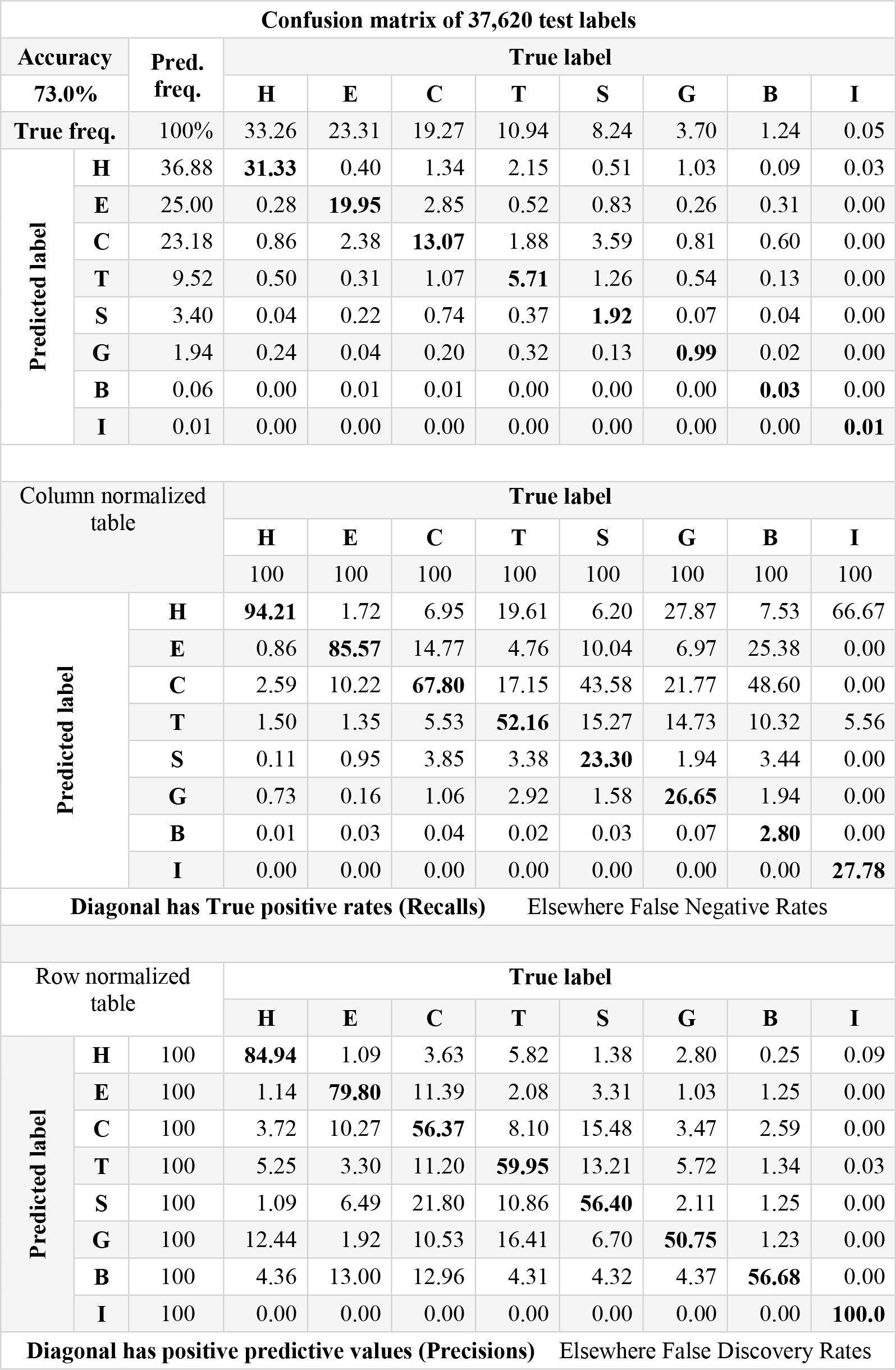
Confusion matrices for 8-label alphabet: H, E, C, T, S, G, B, and I. (1) Confusion matrix, (2) true positive rates (recalls) and false negative rates and (3) positive predictive values (precisions) and false discovery rates of SecNet on Test2018 test set.

**Table 3.**
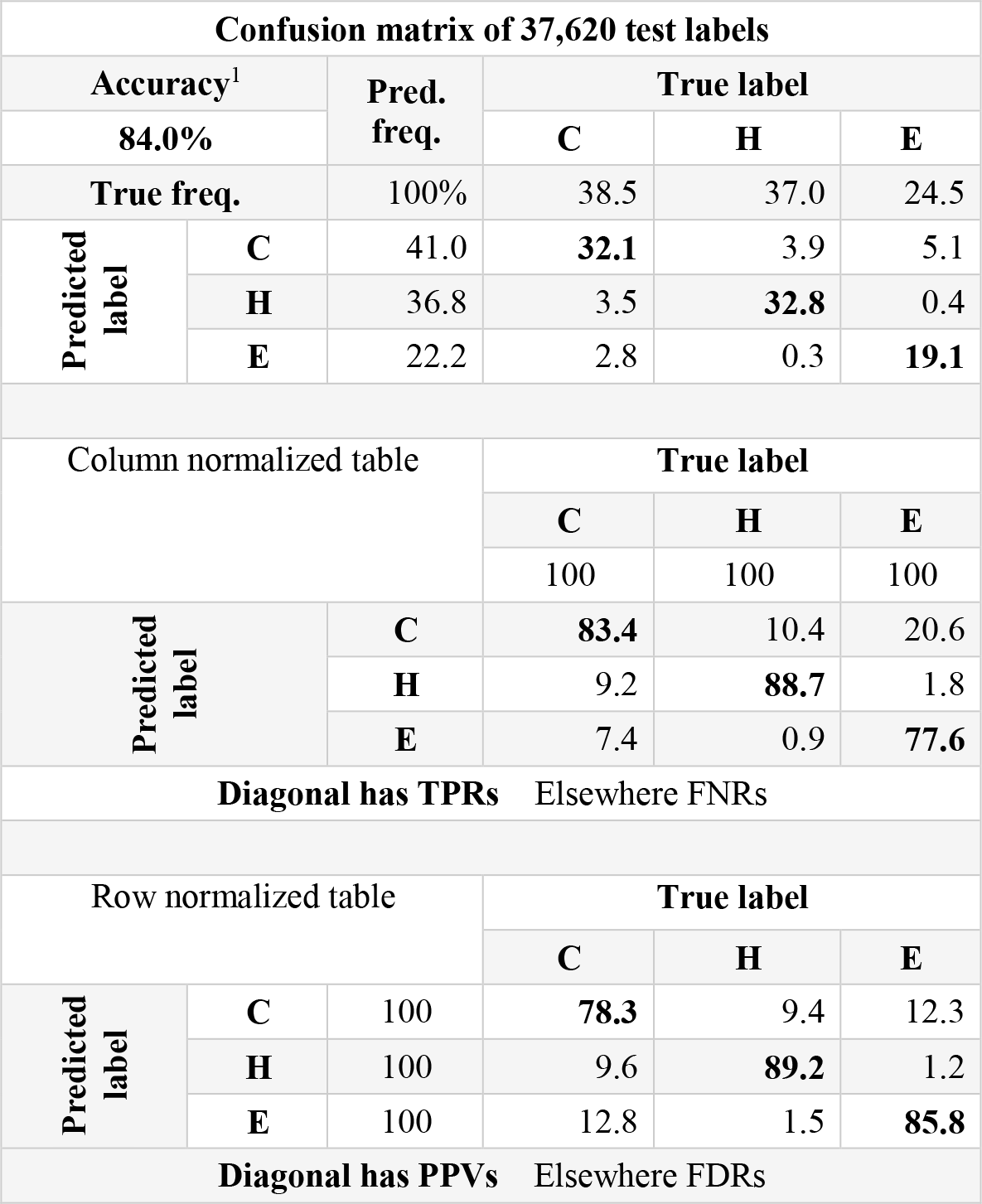
*Rule #1* of the popular 3-label (*H*, *C*, and *E*) alphabet: (1) confusion matrix, (2) true positive rates (recalls) and false negative rates and (3) positive predictive values (precisions) and false discovery rates of *SecNet*. The most common and harder 3-label (*H, C, E*) Rule #1 is defined as (G, I) → H, (B) → E and (B, S) → C. Depending on different studies, it is 2-3% lower in accuracy than easier Rule #2. ^1^The overall accuracy is summation of the main diagonal of the confusion matrix.

We also tested *SecNet* with the popular *CB513* test set derived by Cuff and Barton in 1999 for benchmarking of competing secondary structure prediction software [91] and adopted for accuracy assessment many times [73, 77, 79-81]. *CB513* has several deficiencies. First, disordered residues that have no atoms present in the coordinates are deleted from the sequences and the DSSP strings altogether. This means that sequence profiles and HMMs will be distorted since proteins in the sequence database will seem to have insertions relative to the query protein even when they do not. Second, it has relatively poor average and maximal resolutions (2.11 Å and 3.5 Å) compared to sets that can be derived from the PDB today (e.g., the *Test2018* test set has average and maximal resolutions of 1.83 Å and 2.2 Å). The *CB513* data set protocol initially required ≤ 2.5 Å resolution; however, 5% of its entries have resolution between 2.5 and 3.5 Å resolution. Moreover, 32% of *CB513* entries have free R-factor in the worst 25^th^ percentile for the resolution of the deposited structures in the PDB. Such models raise doubts about the quality of the refined structures [92]. Third, *CB513* has sequences of PDB chains that are fragmented into domains, which results in sequences 2.2 times shorter on average than their full PDB chains. *CB513* has sequences as short as 20 amino-acid residues; 5% and 20% of chains in *CB513* are shorter than 40 and 80 residues respectively. Finally, secondary structure labels were generated with an obsolete version of *DSSP* that was available in 1998.

Nevertheless, because we eliminated proteins from our training and validation sets with more than 25% sequence identity with any chain in *CB513* (Fig 1), we can fairly compare *SecNet* with numerous previous reports on *CB513*. We benchmarked *SecNet* on *CB513* and calculated 8- label, *Rule #1* 3-label, and *Rule #2* 3-label accuracies of 72.3%, 84.3%, and 86.3% respectively (Table 4), which are close to the *Test2018* accuracies with −0.7%, +0.3% and +0.3% differences; a lower value for 8 *DSSP* labels may be related to the *CB513* deficiencies described above.

**Table 4.**
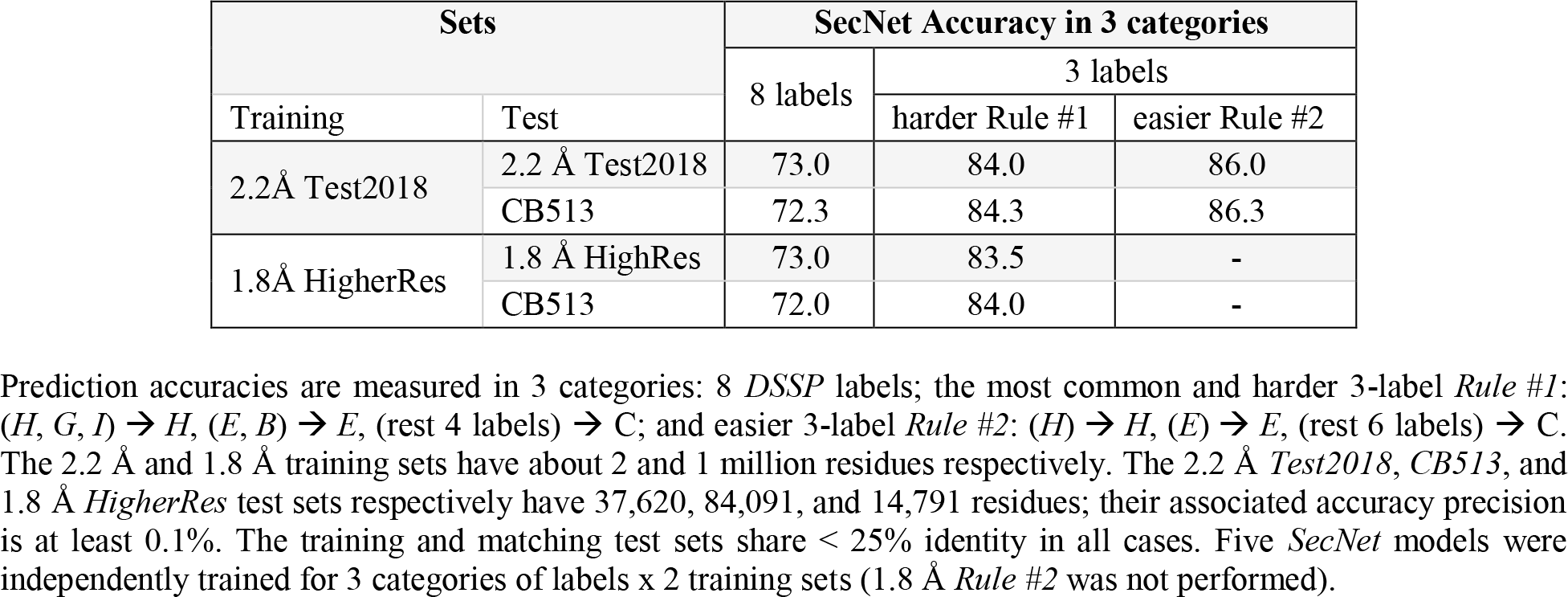
Prediction accuracy of our SecNet software in 3 categories of labels on 3 test sets.

Concerned with discrepancies in X-ray crystal structures with resolution up to 2.2 Å in *Set2018* and up to 3.5 Å in *CB513*, we explored more stringent structural quality parameters that result in smaller training and test sets, but may result in higher prediction accuracy since secondary structures may be more accurately determined especially for 8 labels. To test this, we followed the same protocol to produce training, validation, and test sets with a 1.8 Å cutoff (*‘HigherRes*’ sets), and then retrained the *SecNet* neural network. We then benchmarked *SecNet* on the 1.8 Å *HigherRes* and *CB513* test sets. The 3-label accuracies are lower by 0.3-0.5% for both test sets; 8- label accuracy is the same for the new test set compared to the 2.2 Å test set; it is lower by 0.3% for *CB513* (rows 1-2 vs. 3-4 in Table 4). The modest decreases are likely due to (1) the smaller data set size – 1,224,479 vs. 2,074,301 amino acids in the 1.8 Å and 2.2 Å data sets respectively and (2) a small impact from the ground-truth secondary-structure assignment at the higher resolution of 1.8 Å. Since *SecNet* has approximately the same accuracy on the 2.2 Å *Test2018* as it has on the 1.8 Å *HigherRes*, and since the 2.2 Å training and testing sets are much larger, all further results, discussion, and final benchmarking are reported on *Test2018* at 2.2 Å resolution.

Next, we compared *SecNet* with 12 fifth-generation template-free methods with reported *CB513* accuracies in recent publications (Table 5). Our *SecNet* accuracies are 2-9% higher than 11 out of 12 previous methods with only *eCRRNN* reporting a higher accuracy by 1.7% on 8-label predictions and 1.1% higher on 3-label predictions. We chose to test *eCRRNN* directly as well as *DeepCNF*, both having the highest reported accuracies on *CB513*. *DeepCNF* uses a CNN with 5 layers and conditional random field as an additional layer and as input includes 42 features for each residue: 21 from a PSSM profile and 21 from one-hot encoding. *DeepCNF* has an effective window size of 51 amino acids. The *eCRRNN* architecture combines 6 blocks as a mixture of convolutional, residual and bidirectional recurrent NNs. The input to *eCRRNN* consists of 50 features for each residue – 20 values from a PSSM profile, 7 physical properties, a conservation score, and a set of twenty-two 22-dimensional orthogonal vectors. The second-block BRNNs process the full amino-acid sequence by reading all residues on the left and right from the residue subject to prediction, making the window width effectively unlimited.

**Table 5.** CB513 accuracy of the 5^th^-generation template-free secondary-structure prediction methods.

| Method | Seq id of *CB513* and training data | 8 labels |  |  | Harder 3-label Rule #1 |  |  | Easier 3-label Rule #2 |  |  |
| --- | --- | --- | --- | --- | --- | --- | --- | --- | --- | --- |
|  |  | accuracy | source | diff | accuracy | source | diff | accuracy | source | diff |
| Our *SecNet* (2019) | < 25% | 72.3 | authors | 0.0 | 84.3 | authors | 0.0 | 86.3 | authors | 0.0 |
| **eCRRNN (2018)** | unspec | **74.0→70.2** | auth→calc | -2.1 | 81.2 | calculated | -3.1 | **87.3→84.3** | auth →calc | -2.0 |
| MUFOLD-SS (2018) | unspec | 70.6 | authors | -1.7 | - | - | - | 82.7 | eCRRNN | -3.6 |
| CNNH_PSS (2018) | < 25% | 70.3 | authors | -2.0 | - | - | - | - | - | - |
| DCRNN (2016) | < 25% | 69.7 | authors | -2.3 | - | - | - | 84.0 | authors | -2.3 |
| DCNN (2017) | < 25% | 70.0 | authors | -2.6 | - | - | - | - | - | - |
| **DeepCNF (2016)** | < 25% | **68.3→69.1** | auth→calc | -3.2 | **82.3→81.8** | auth→calc | -2.5 | - | - | - |
| BLSTM (2015) | < 25% | 67.4 | authors | -4.9 | - | - | - | - | - | - |
| GSN (2014) | < 30% | 66.4 | authors | -5.9 | - | - | - | - | - | - |
| SSpro, free (2014) | Any | 63.5 | DeepCNF | -8.8 | 78.5 | DeepCNF | -5.8 | - | - | - |
| JPRED4 (2015) | Any | - | - | - | 81.7 | DeepCNF | -2.6 | 79.6 | FSVM | -6.7 |
| FSVM, free (2018) | < 25% | - | - | - | - | - | - | 82.9 | authors | -3.4 |
| SPIDER2 (2015) | Any | - | - | - | - | - | - | 81.2 | FSVM | -5.1 |
| PSIPRED (1999) | Any | - | - | - | 79.2 | DeepCNF | -5.1 | - | - | - |

*CB513* accuracies were compiled from recent papers of the fifth-generation, template-free methods. Accuracy taken from an original publication has “authors” tag. If a study did not utilize the *CB513* test set but a third-party benchmarked the method, the latter source is reported instead and the sequence identity to *CB513* is listed as “Any.” Some “authors” papers did not specify an identity cutoff between their training set and their test sets, including *CB513* (“unspec”). The widely used *PSIPRED* program belongs to the third generation. We recalculated the *eCRRNN* and *DeepCNF* accuracies (“calc.”) with the original *CB513* (indicated after the →).

Since *DeepCNF* and *eCRRNN* were submitted for publication prior to the end of 2018, our *Test2018* set is independent of their training sets (at 25% sequence identity). We applied both methods to the *Test2018* test sequences, and determined that our *SecNet* has higher accuracies than *DeepCNF* by 3.3-3.4% (Table 6). Our method’s accuracies are 2.0-2.2% higher than *eCRRNN* on *Test2018*. Next, we ran both of these methods on the original *CB513* test set. The *DeepCNF* accuracies that we calculated on *CB513* closely reproduce those of the authors with slightly higher 8-label and slightly lower 3-label accuracies (Table 5).

**Table 6.** Benchmarking of SecNet, DeepCNF, and eCRRNN on Test2018.

| Test set | Method | 8 labels |  | 3 labels |  |  |  |
| --- | --- | --- | --- | --- | --- | --- | --- |
|  |  |  |  | harder Rule #1 |  | easier Rule #2 |  |
|  |  | accur | diff | accur | diff | accur | diff |
| Test2018 | **SecNet** | **73.0** | **0.0** | **84.0** | **0.0** | **86.0** | **0.0** |
|  | eCRRNN | 70.8 | -2.2 | 81.8 | -2.2 | 84.0 | -2.0 |
|  | DeepCNF | 69.6 | -3.4 | 80.7 | -3.3 | - | - |

**Table 7.** Evidence of prediction accuracy overestimation of *SecNet* on *Set2018* validation and training sets relative to the test set.

|  | Set2018 | Accuracy |  |
| --- | --- | --- | --- |
|  |  | SecNet | Diff. from Test |
| 3 labels, Rule #1 | Test | 84.0 | 0.0 |
|  | Validation | 86.1 | +2.1 |
|  | Favorable Validation | 88.0 | +4.1 |
|  | Training | 88.3 | +4.3 |
| 8 labels | Test | 73.0 | 0.0 |
|  | Validation | 75.5 | +2.5 |
|  | Favorable Validation | 77.8 | +4.8 |
|  | Training | 79.8 | +6.8 |

The test accuracy was probed only once for reporting accuracy results. Accuracy probes on the validation set were done multiple times for design decisions (~10s of times), hyper-parameter tuning (~100s), saving the best model during training (~100 probes per training x ~1,000s trainings = ~10,000s), and rejecting more complex *SecRes* residual NN (~10s). The “favorable validation” accuracy was achieved by splitting the whole training set with a 1:9 ratio into validation and reduced training sets more than 10 times (~100s). The models were individually trained, and the validation set with the best accuracy was chosen. The training set accuracy was probed few million times per training (~1,000,000s) during gradient-descent optimization. It is considerably below 100% owing to the regularization technique with 30% dropout. The overestimated numbers underline the importance of having separate test, validation, and training sets and probing the test set only once for reporting unbiased results.

When the authors tested *DeepCNF* on *CB513*, they excluded 31 shorter entries, and this may account for these small observed differences. In contrast, our recalculated *CB513* accuracies for *eCRRNN* indicated that the authors overestimated their accuracy by 3.0-3.8% (Table 6). We excluded the possibility that we incorrectly executed the *eCRRNN* software by reproducing their results for the three tests sets included with their software. With these findings we updated the *CB513* accuracy for *eCRRNN* in Table 5 with our calculated values. With these accuracy revisions, *SecNet* is more accurate than *eCRRNN* by 2.0-2.2% on *Test2018* and 2.0-3.1% on *CB513*, depending on the label set.

Several programs (including *eCRRNN*) in Table 5 do not specify a similarity cutoff (“unspec”) between training and test sets. Some apply a higher value [82, 100] than the commonly used 25%, and one program, *PSRSM* [83], did not mention any similarity exclusion technique between these sets in their publication. When training set proteins similar to the testing set are not excluded properly, it is likely that such methods overestimate their accuracy by overtraining. Accuracy may also be overestimated when a single data set is used as both validation set and testing set. To demonstrate this, we compared the accuracy on our testing set and our validation set achieved during training. As expected, the accuracy on the validation set is higher than the accuracy on the test set by 2.5% (75.5% vs. 73.0%) for 8-labels and 2.1% (86.1% vs. 84.0%) for *Rule #1* (Table 7). The accuracy on the training set is 6.8% higher (79.8% vs. 73.0%) for 8 labels and 4.3% higher (88.3% vs. 84.0%) for *Rule #1* 3-label predictions. The striking over-estimation for the validation and training sets demonstrates the importance of separate training, validation, and testing data sets. As another test, if we randomly permute validation and training splits many times with the 1:9 ratio, individually train each, and select the best model based on the highest validation accuracy, we gain additional 2.3% and 2.0% improvements in the 8-label and *Rule #1* 3-label validation accuracy with no improvement in the test accuracy (Table 7). In this case, the validation accuracy is boosted due to a favorable random re-distribution of proteins in the training and validation sets.

Finally, the accuracy of secondary structure prediction methods for 3 labels depends on the helix, sheet, and coil fractions in the test set. If we oversample *E* or *H*, we can vary the accuracy from 78% to 89% accuracy respectively (Fig 3). As it turns out, our training and test sets and *CB513* are very similar to each other in terms of the 3-label fractions of both *Rule #1* with 37%, 24%, and 39% and *Rule #2* with 34%, 23%, and 43% for helix, sheet, and coil respectively (Fig 3 and Table 3). This should be kept in mind when deriving new test sets.

**Fig 3.**
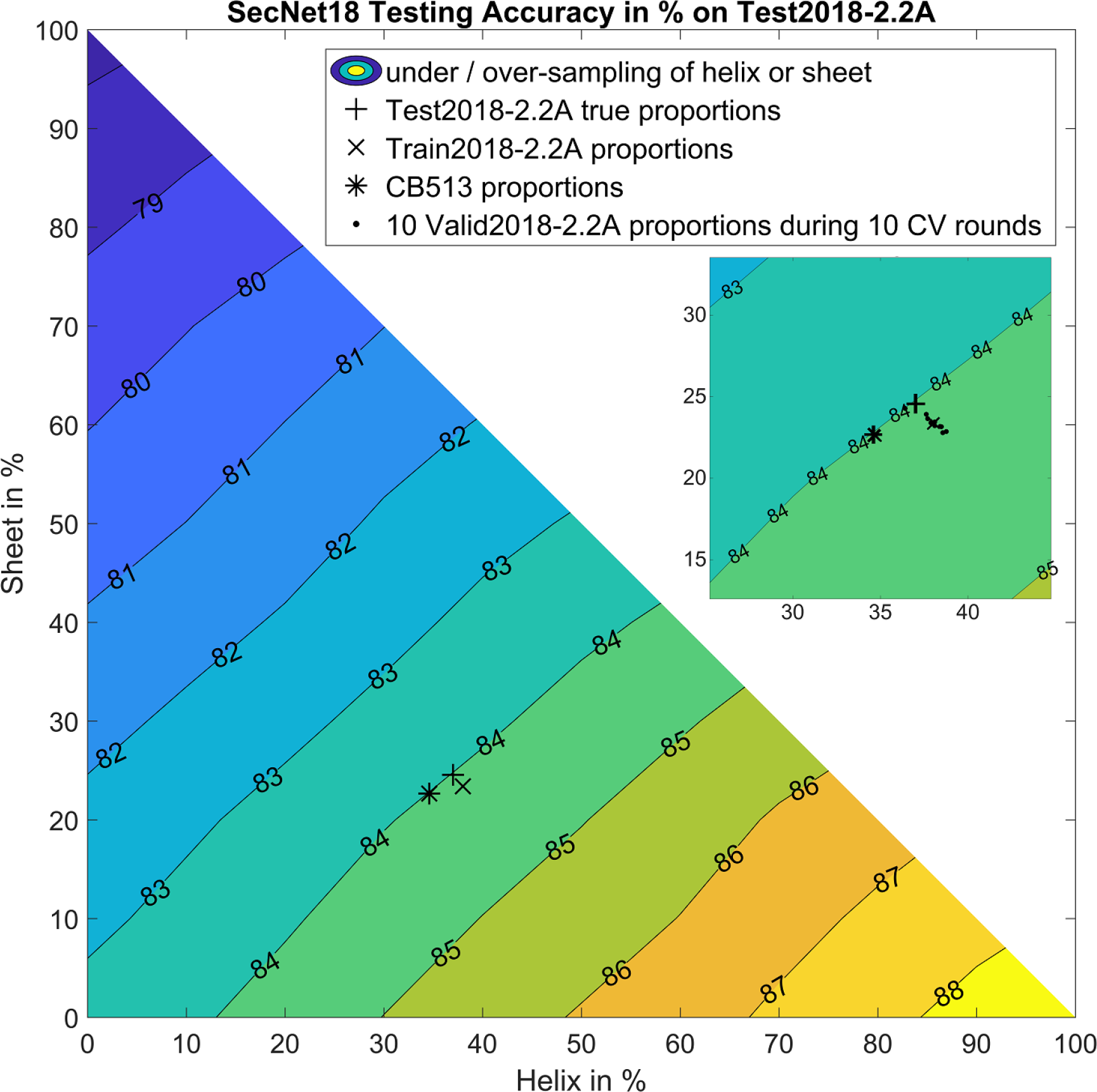
Variation of 3-label *Rule #1* accuracy as a function of proportions of helix (*H%*), sheet (*E%*) and coil (*C%* = 100% – *H%* – *E%*) in *Test2018* test set. The *Rule #1* accuracy of *SecNet* on *Test2018* test set is 84.0% and shown with a black plus. The contour plot demonstrates how *SecNet* prediction accuracy is skewed when a test set is enriched with helix. For example, if the underlying test set has 100% helices, the accuracy is 88.7% (bottom right). If it is 100% sheet, the accuracy is 77.6% (top left). If it is 100% coil, it is 83.4% (bottom left). The *SecNet* accuracies are shown for the label proportions of *CB513* test set (black star), training + validation sets (black cross) and 10 validation sets, one from each round of cross validation (10 black dots).

#### Choices that affect the accuracy of secondary structure prediction

We explored many different ideas and options during the development of *SecNet*. We can divide these choices into three categories: (1) neural network type, architecture, and complexity; (2) input features; and (3) how to perform training of the neural network. After optimizing *SecNet* and using the *Test2018* set only once (Table 4), we performed an ablation study of the effect of these factors on the 8-label accuracy. Some of the necessary CNNs were produced during the development of *SecNet* and some needed to be generated for the ablation study. The results are shown in Fig 4. The first line of the figure lists the three accuracy measures achieved by making various choices shown in the rest of the figure: 86.0% for the 3-label accuracy from *Rule #2* (*H*→*H*; *E*→*E*; all others →*C*), 84.0% for the 3-label accuracy from *Rule #1* (*G*, *H*, *I*→*H*; *B*, *E*→*E*; all others *C*), and 73.0% for the 8-label accuracy. Starting from the middle of the second line, using a “larger HMM DB”, “4 days of training instead of 1”, and “ensemble of 10 models” raise the accuracy from 72.0% to 73.0% in increments of 0.1%, 0.3%, 0.6% respectively. A majority of individual accuracy improvements in the figure are a fraction of a percent; it is the cumulative effect of many choices that makes a significant improvement in accuracy.

**Fig 4.**
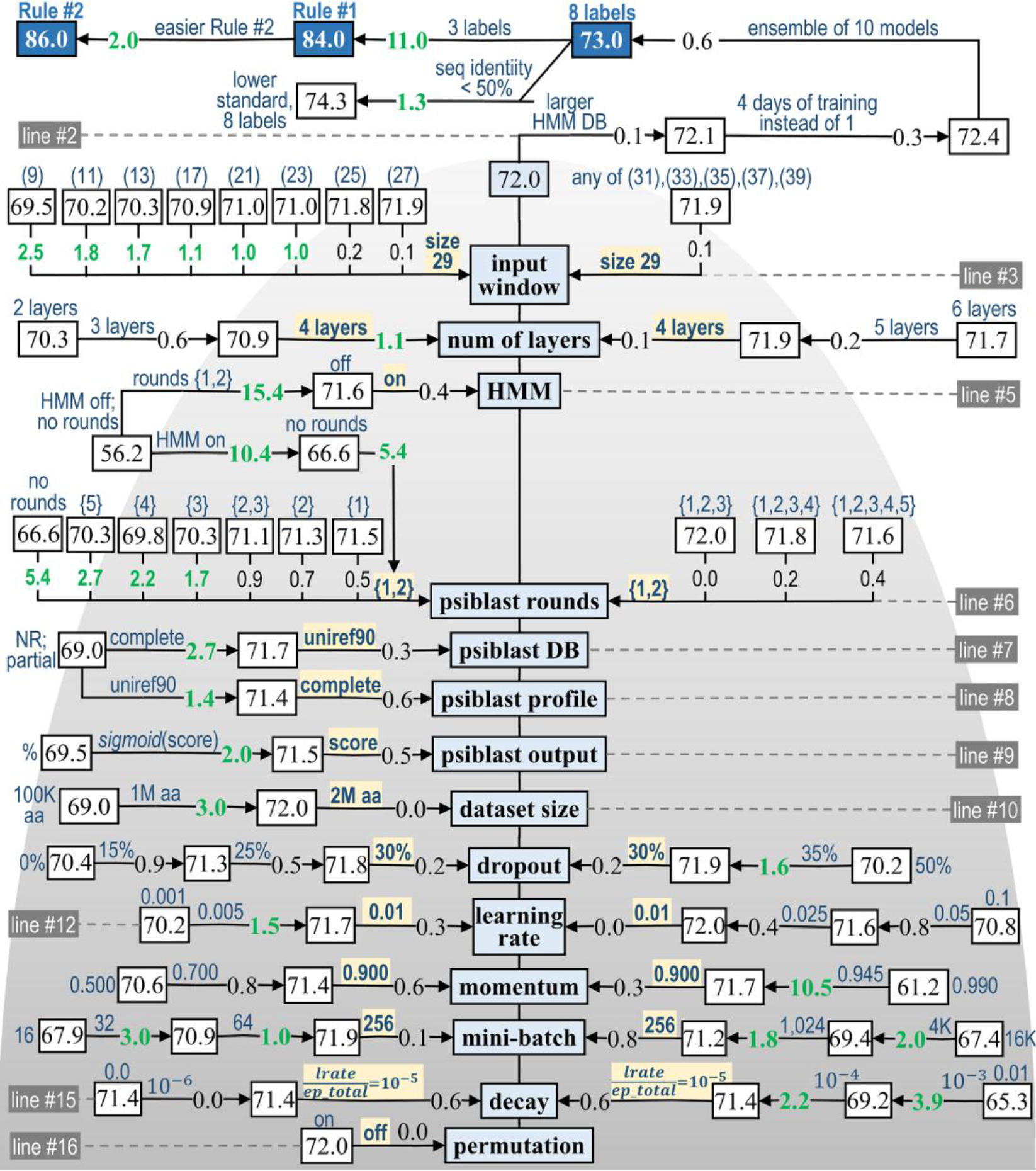
Ablation study of *Test2018* accuracy of *SecNet*. Accuracy is presented as a function of factors (blue boxes in the middle) from 3 groups: 1) NN architecture and complexity (top); 2) input features and databases (center); and 3) hyper-parameters of the training pipeline (bottom). The test accuracies displayed in black uncover unbiased estimates for publication purposes *after* all choices of the final model parameters were made; the validation set was used to make these choices. *Lines 1-2*: Actions such as “4-day training” or “ensemble of 10 cross-validated models” led to further improvement. *Line 1*: The accuracy increases with switch of 8→3 labels by replacing a harder *Rule #1* to easier *Rule #2* and with a weaker 25%→50% maximal sequence identity between the test and training sets. Each arrow indicates a direction of favorable parameter change and embeds associated accuracy gain. Parameter values are in blue. The optimal parameter values are highlighted in yellow and shown in the middle with smaller and larger values on the left and right. The cumulative effect from multiple actions is a sum of individual accuracy increments (in black). Stronger effects (≥1%) are shown in green.

Each line below the second line ablates one of the factors, such as a feature, algorithmic, or training choices described in a blue rectangular box; the highest accuracy of 72.0% is achieved with a set of the optimal parameters shown in the middle of the figure. Non-optimal parameters are ordered on the left and right with lower and higher values compared to the optimal values respectively.

The first category of choices involves the architecture of the neural network and its complexity. For example, secondary structure formation is influenced by short, middle, and long- range inter-residue interactions. We were interested to see how the accuracy relates to the size of the sequence “input window” (line 3 in Fig 4) centered on a residue subject to prediction. To investigate this, we varied the input window size, and observed increasing accuracy from 69.5 to 71.9% for a window of 9 to 27 (left), the highest 72.0% for the optimal window of 29 (middle) and subsequent degradation to 71.9% for a window of 31 to 39 (right). The severe degradation of the accuracy (1.7-2.5%) was observed for 9-13 amino-acid windows, moderate reduction (0.1- 1.1%) was for 17-27 windows and very minor decrease (0.1%) was for 31-39 windows.

Another important factor in secondary structure prediction accuracy is the kind of neural network and the number of layers. The 4-layer NN is more accurate by 1.7% and 1.1% than 3- layer and 2-layer NNs and by 0.1% and 0.3% than 5-layer and 6-layer NNs (line 4). Wang et al. made a similar observation of accuracy saturation at 5-7 layers [79]. We also tested a more complex residual neural network with 20-40 layers and a 21-51 amino-acid window size, and observed the same or worse accuracy than *SecNet*’s traditional CNN with 4 layers (Fig A in S1 Supporting information). Coupled with the results from Table 5, we conclude that a CNN is adequate for secondary structure prediction, despite availability of newer and more complex neural architectures.

The second category of choices consists of the features used as NN inputs. The use of sequence profiles, as demonstrated by many programs over the last 30 years, has a strong positive impact on accuracy (line 5). If both the *PsiBlast* and *HMM* features are disabled, the accuracy drops from 72.0% to only 56.2% (“HMM off; no rounds”), contributing to the largest observed impact of all factors we explored in the ablation study. When either *HMM* or *PsiBlast* are individually enabled, the accuracy jumps to 66.6% and 71.6% respectively.

A choice of database for the *PsiBlast* searches is not that critical. *Uniref90* is only 0.3% better than the *NR* non-redundant sequence database from NCBI (line 7), probably because for some targets *NR* produces too many nearly identical hits, and subsequently skews the *PsiBlast* profile too much in their favor. There is a *PsiBlast* command-line argument that determines how many of the top hits are included in a profile; if the profile is limited to the top 15 hits, the accuracy suffers by a hefty 2.7% for *NR* (“NR; partial” vs. *NR*’s “complete” in line 7) and only 0.6% for *Uniref90* (line 8). *PsiBlast* produces profiles both in log-odds form and amino-acid percentages. The log-odds form (“score”) is better than the percentage values by 2.5% (line 9). It is common to use a sigmoid function to transform a feature into a 0 to 1 range. The log-odds score is also better than their sigmoid function values (“*sigmoid*(score)”)—by 0.5%.

*PsiBlast* profiles generated after a certain number of rounds have a significant effect (line 6). Profiles obtained from the third, fourth, and fifth rounds contain homologues that are very distant from a target sequence and therefore may contain significant differences in secondary structure, which seem to adversely affect the accuracy. The accuracy of neural networks trained on profiles after the third, fourth, and fifth rounds alone drops respectively by 1.7%, 2.2%, and 2.7% relative to the best option which includes both profiles after the first and second rounds. This best option is higher by 0.5% and 0.7% than using the first or second round profiles alone. We observe a steady but modest degradation of accuracy by 0.0% or 0.2% or 0.4% when a combination of rounds 1-3, 1-4, or 1-5 are used. A possible explanation is that the additional rounds decrease the data/parameter ratio. The round 2-3 combination is worse by 0.9% than the round 1-2 combination.

The third category of choices involves the training options. A difference in training data set size of 1 or 2 million amino-acids has no effect, but a 10-fold training data set reduction to 100 thousand amino acids pushes accuracy down by 3% (line 10). For regularization, we chose the dropout approach by randomly forgetting 30% of the model parameters during each training iteration (line 11). This approach results in 1.6% higher accuracy compared to training without any dropout. The 30% dropout is optimal; the accuracy drops when dropout is either decreased or increased from 30%. CNN is relatively insensitive to small changes in learning rate from 0.01 and momentum from 0.900. However, larger deviations from these values result in significant accuracy losses; for example, the accuracy loses 0.3-1.8% and 0.4-1.2% when the learning rate is 0.001-0.005 and 0.05-0.1 respectively (line 12). A momentum of 0.500 instead of 0.900 results in a 1.4% drop while a value of 0.990 reduces accuracy by 10.8% (line 13).

Each weight update in training our network involves calculating derivatives of the loss function for a batch of data, called a “mini-batch.” If the mini-batch is too small, then the calculated derivatives will be too noisy; mini-batch sizes of 16-64 instead of 256 amino acids produced losses of 0.1-4.1% in accuracy. A mini-batch size that is too large may produce too stable loss function gradient and may result in premature convergence of the model to a less optimal set of parameters; mini-batch sizes from 1,024 to 16,384 resulted in a loss of 0.8-4.6%. (line 14).

In order to find a better optimum during the training and especially toward its end, it is critical to reduce the size of gradient-descent steps by decreasing the current learning rate (“decay”). A formula-based decay of “learning rate / epoch total” which is equal to 10^-5^ with the optimal learning rate of 0.01 and 650 epochs for 4 days, is better by 0.6% than no decay (0.0) or smaller decay of 10^-6^; it is also better by 0.6, 2.8 and 6.7% than a larger decay of 10^-4^ or 10^-3^ or 1.1 respectively (line 15). Finally, if we permute the data set and split into the training and validation sets *many* times, the validation accuracy becomes overestimated by 2.3% due to easier targets in the favorable random validation set; however, we observe no improvement (0.0%) in the testing accuracy (line 16). Other training options were described earlier (lines 1-2), resulting in a final accuracy of 73.0%.

We conclude that the best results are achieved with good choices for input features, sequence databases and arguments, and the training pipeline and regularization techniques. It appears that the NN type, architecture and complexity, and the associated number of training parameters are not as critical as long as the NN is not too simple.

#### Practicality of secondary structure prediction: which and how many labels

Most existing secondary structure prediction programs have been trained and tested on either 8-label or 3-label data sets. In most cases, the 8 *DSSP* labels are reduced to 3 by treating 3_10_ helices (“G” in *DSSP*) and π helices (“I” in *DSSP*) as *H*, and single-residue beta strands (beta bridges or “B” in *DSSP*) as *E*; the other labels (“C” or coil, “T” or turns, “S” or bends) are reduced to *C*. We wondered how label sets with more than 3 but fewer than 8 labels would behave and how these might be developed.

To investigate this, we calculated the confusion matrix for 8-label *SecNet* (Table 2) and both the column and row-normalized versions of the confusion matrix. The column-normalized table (Table 2, middle) provides true positive rates (“recalls”) along the main diagonal (TPR=TP/(TP+FN)) and false negative rates (FNR=FN/(TP+FN)) off the diagonal; the row-normalized table (Table 2, bottom) reports positive predictive values (“precisions”) along the diagonal (PPV=TP/(TP+FP)) and false discovery rates off the diagonal (FDR=FP/(TP+FP)).

The column-normalized confusion matrix shows that only labels *H*, *E*, *C*, and *T* have true positive rates over 50%. The TPR for beta bridges (“B”) is only 2.8%. Experimental *G* and *I* are predicted as *H* more often than their true labels. Bends (“S”) are predicted as *C* almost twice as often as they are predicted as *S*. The row-normalized confusion matrix shows that all 8 labels have positive predictive values over 50%, but they are more than 60% for only *H* (85%), *E* (80%), and *I* (100%). *I* labels, however, are only 0.05% of the true labels and 0.01% of the predictions (Table 2, top).

From a practical point of view, predicted secondary structure labels are useful if they are accurate (high PPV and TPR), commonly observed, and lead to effective sampling strategies in tertiary structure prediction, which might be performed though the use of fragments or from dihedral angle distributions. From this point of view, the *S* and *B* labels are not very useful. The *S* label indicates a “bend” when the angle Cα(i-2) – Cα(i) – Cα(i+2) is less than 110° and residue *i* is not *H*, *B*, *E*, *G*, *I*, or *T* [2]. The *B* label indicates a beta bridge, which is simply a one-residue backbone-backbone hydrogen bond, resembling a one-amino-acid beta-sheet strand. Since *S* and *B* have low TPR values and are impractical to sample effectively, it makes sense to convert them to the catch-all label, *C,* and sample them from Ramachandran distributions generated from residues not in helices or sheets [101]. Similarly, the *I* label is so rare that it may be converted to *H*, since about 75% of the time it occurs as the first or last turn in an alpha helix. Thus, after these conversions, we are left with *H*, *E*, *C*, *T*, and *G*.

We trained and tested a 5-label version of *SecNet* (*H*, *E*, *C*, *G*, *T*), and achieved an accuracy rate of 78.3% (Table 8), compared to 84.0% for the *Rule #1* 3-label predictions, 86.0% for the *Rule #2* 3-label predictions, and 73.0% for 8-label predictions. The TPR values for the 5-label rules for *H*, *C*, *E*, *T*, and *G* are 94%, 78%, 84%, 52%, and 34% respectively. The PPV values are 89%, 73%, 84%, 69%, and 55%.

**Table 8.**
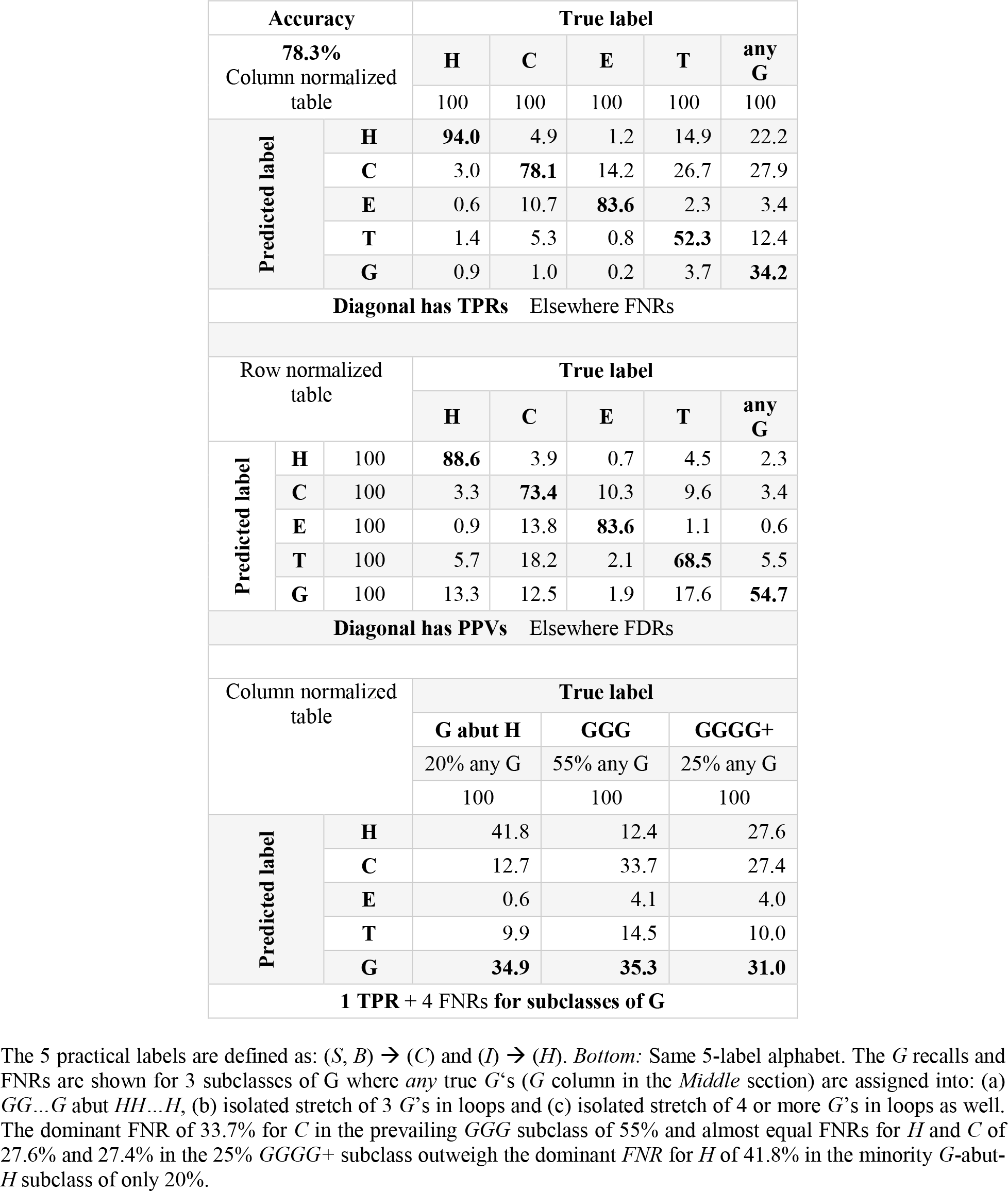
5-label practical alphabet: H, C, E, T, and G: (1) true positive rates (recalls) and false negative rates and (2) positive predictive values (precisions) and false discovery rates of SecNet, (3) TPRs and FNRs for three subclasses of G: (a) G abut HH…H, (b) isolated GGG in loops and (c) isolated GGGG+ in loops.

The label *G* (3_10_ helices) remains problematic in the 5-label scheme with false negative rate of 28%, 22%, and 12% for *C*, *H*, and *T* compared to a 34% true positive rate for *G*. In a recent study on beta turns [102], we divided 3_10_ helices into three categories: 3_10_ helices that abut alpha helices (20% of *G*), isolated 3-residue 3_10_ helices (the most common length by far) in loop regions and not abutting alpha helices (55% of *G*), and isolated 3_10_ helices of length 4 or more (25% of *G*). 3_10_ helices that abut alpha helices might be viewed as a distortion of the alpha helix and therefore might be more similar to *H* labels. Isolated 3_10_ helices of length 3 strongly resemble the succession of two Type I beta turns (with distorted backbone dihedral angles for the first turn compared to other Type I turns) [102].

We analyzed the TPR and FNR values for these three categories of residues in 310 helices from the 5-label predictions (bottom of Table 8). With *SecRes* trained on a 5-letter scheme, *G* residue stretches that abut alpha helices have a 35% true positive rate and false negative rates of 42%, 13%, and 10% for *H*, *C*, and *T* respectively. Isolated *GGG* segments in loops have a 35% true positive rate and false negative rates of 12%, 34%, and 15% for *H*, *C*, and *T* respectively. Finally, isolated 310 helices of length 4 or longer produce a TPR of 31% and FNR values of 28%, 27%, and 10% for *H*, *C*, and *T* respectively. With these results in hand we derive a 4-letter scheme (*H*, *E*, *C*, *T*) by converting all *G* to *C*, since 80% of *G* residues are isolated 3_10_ helices of length 3 or longer. Since the majority of residues in 3_10_ helices come from isolated length-3 helices which are very similar to beta turns, retaining the prediction of turns (label *T*) may enable the sampling of these structures in loop regions. As before, *B* and *S* are converted to *C*, and *I* is converted to *H*. The accuracy for this 4-label practical scheme is 79.5%; the column- and row-normalized confusion matrices for this scheme are provided in Table 9. It achieves very high TPRs of 94%, 79%, and 83% and PPVs of 90%, 75%, and 85% for *H*, *C* and *E*; it achieves satisfactory TPR of 50% and PPV of 71% for *T*. If simpler, practical labels are required, *T* may be replaced with *C* as the best candidate both structurally and in terms of the highest false negative rate of 33% and highest false discovery rate of 22%. The final accuracy for this 3-label prediction scheme is 86.0%; its TPR (recall), FNR, PPV (precision) and FNR are in Table 10.

**Table 9.**
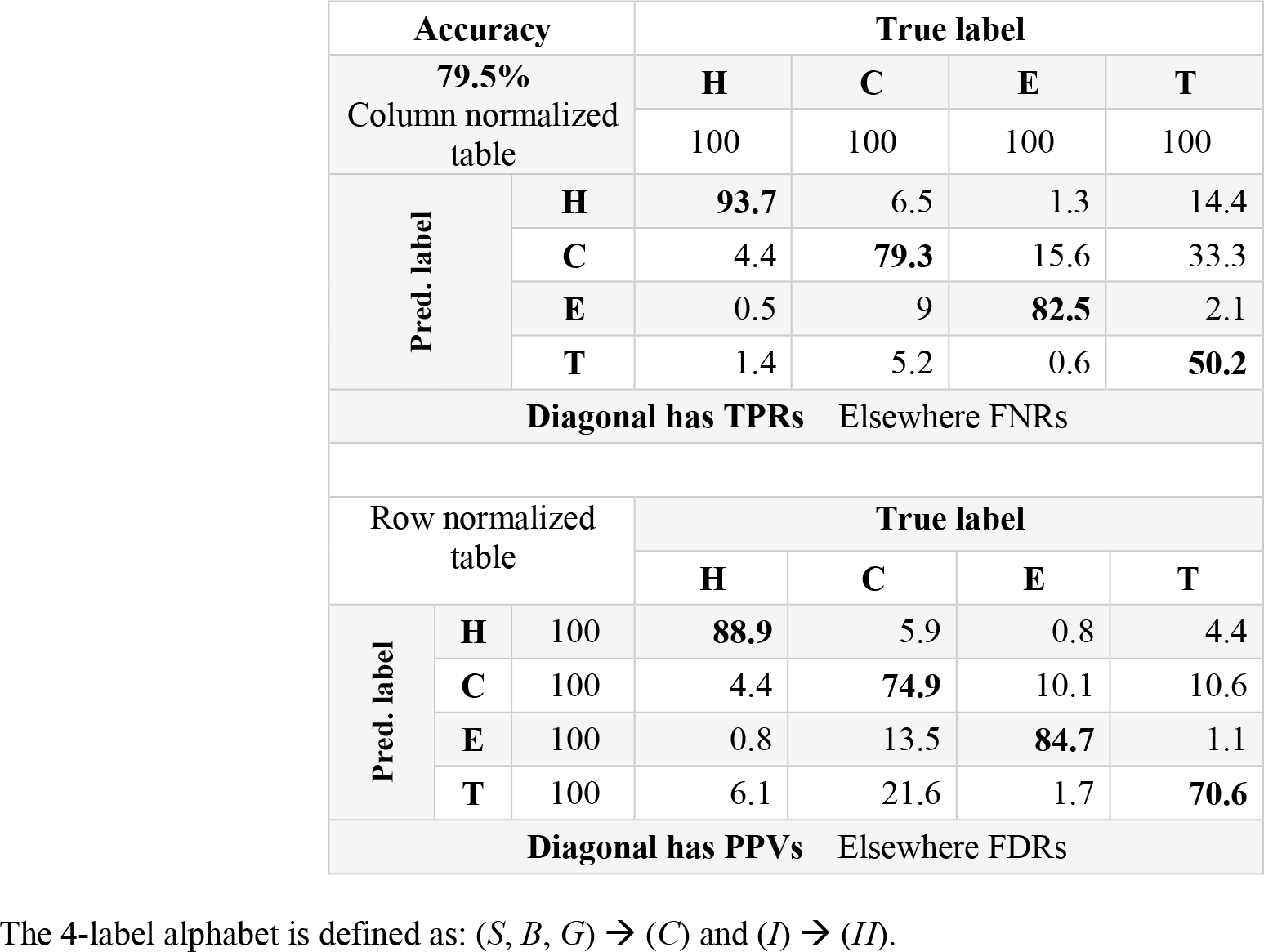
4-label practical alphabet: H, C, E, and T: (1) true positive rates (recalls) and false negative rates and (2) positive predictive values (precisions) and false discovery rates of SecNet.

**Table 10.**
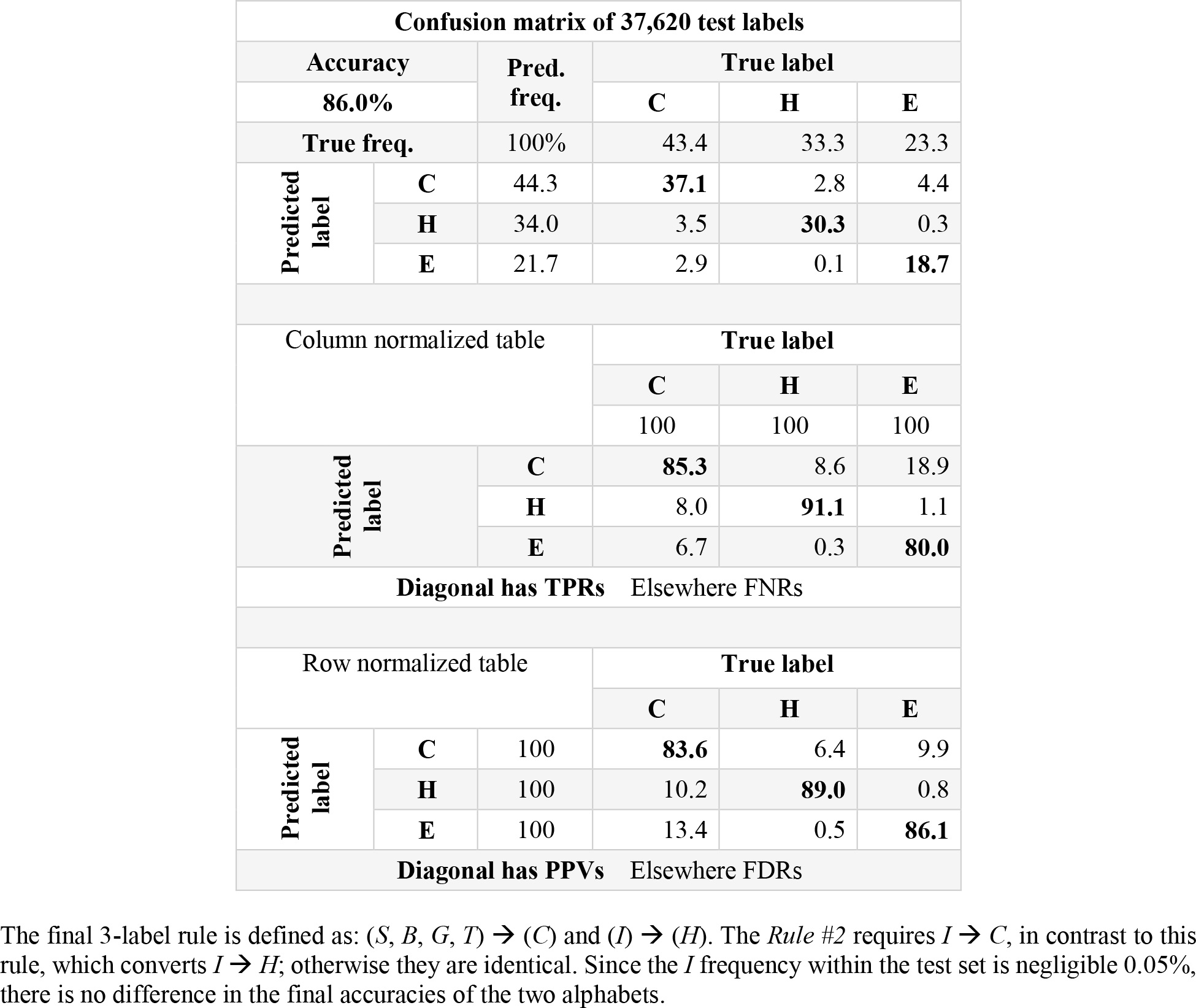
Final 3-label alphabet: H, C, and E: (1) confusion matrix, (2) TPR and TNR and (3) PPV and FDR of SecNet.

| Confusion matrix of 37,620 test labels |  |  |  |  |  |
| --- | --- | --- | --- | --- | --- |
| Accuracy |  | Pred. freq. | True label |  |  |
| 86.0% |  |  | C | H | E |
| True freq. |  | 100% | 43.4 | 33.3 | 23.3 |
| Predicted label | C | 44.3 | **37.1** | 2.8 | 4.4 |
|  | H | 34.0 | 3.5 | **30.3** | 0.3 |
|  | E | 21.7 | 2.9 | 0.1 | **18.7** |
| Column normalized table |  |  | True label |  |  |
|  |  |  | C | H | E |
|  |  |  | 100 | 100 | 100 |
| Predicted label |  | C | **85.3** | 8.6 | 18.9 |
|  |  | H | 8.0 | **91.1** | 1.1 |
|  |  | E | 6.7 | 0.3 | **80.0** |
| Diagonal has TPRs Elsewhere FNRs |  |  |  |  |  |
| Row normalized table |  |  | True label |  |  |
|  |  |  | C | H | E |
| Predicted label | C | 100 | **83.6** | 6.4 | 9.9 |
|  | H | 100 | 10.2 | **89.0** | 0.8 |
|  | E | 100 | 13.4 | 0.5 | **86.1** |
| Diagonal has PPVs Elsewhere FDRs |  |  |  |  |  |

The final 3-label rule is defined as: $(S, B, G, T) \rightarrow (C)$ and $(I) \rightarrow (H)$ . The *Rule #2* requires $I \rightarrow C$ , in contrast to this rule, which converts $I \rightarrow H$ ; otherwise they are identical. Since the *I* frequency within the test set is negligible 0.05%, there is no difference in the final accuracies of the two alphabets.

To summarize, we suggest 5-, 4-, and 3-label schemes as follows:

- 5 labels: (*E*) → *E*, (*H*, *I*) → *H*, (*C*, *S*, *B*) → *C*, (*G*) → *G*, (*T*) → *T* (Table 8)
- 4 labels: (*E*) → *E*, (*H*, *I*) → *H*, (*C*, *S*, *B*, *G*) → *C*, (*T*) → *T* (Table 9)
- 3 labels: (*E*) → *E*, (*H*, *I*) → *H*, (*C*, *S*, *B*, *G*, *T*) → *C* (Table 10)

This last 3-label alphabet is neither *Rule #1* nor *Rule #2* 3-label alphabets found in the literature. It closely resembles *Rule #2* of the 3-label alphabet except for conversion of *I* (0.02% of all *Set2018* residues); due to the negligible frequency of *I*, there is no observable difference in the overall accuracy (86.0%) between this rule and *Rule #2*. In Table 11 we summarize existing and new definitions and compare accuracy of all six alphabets.

**Table 11.**
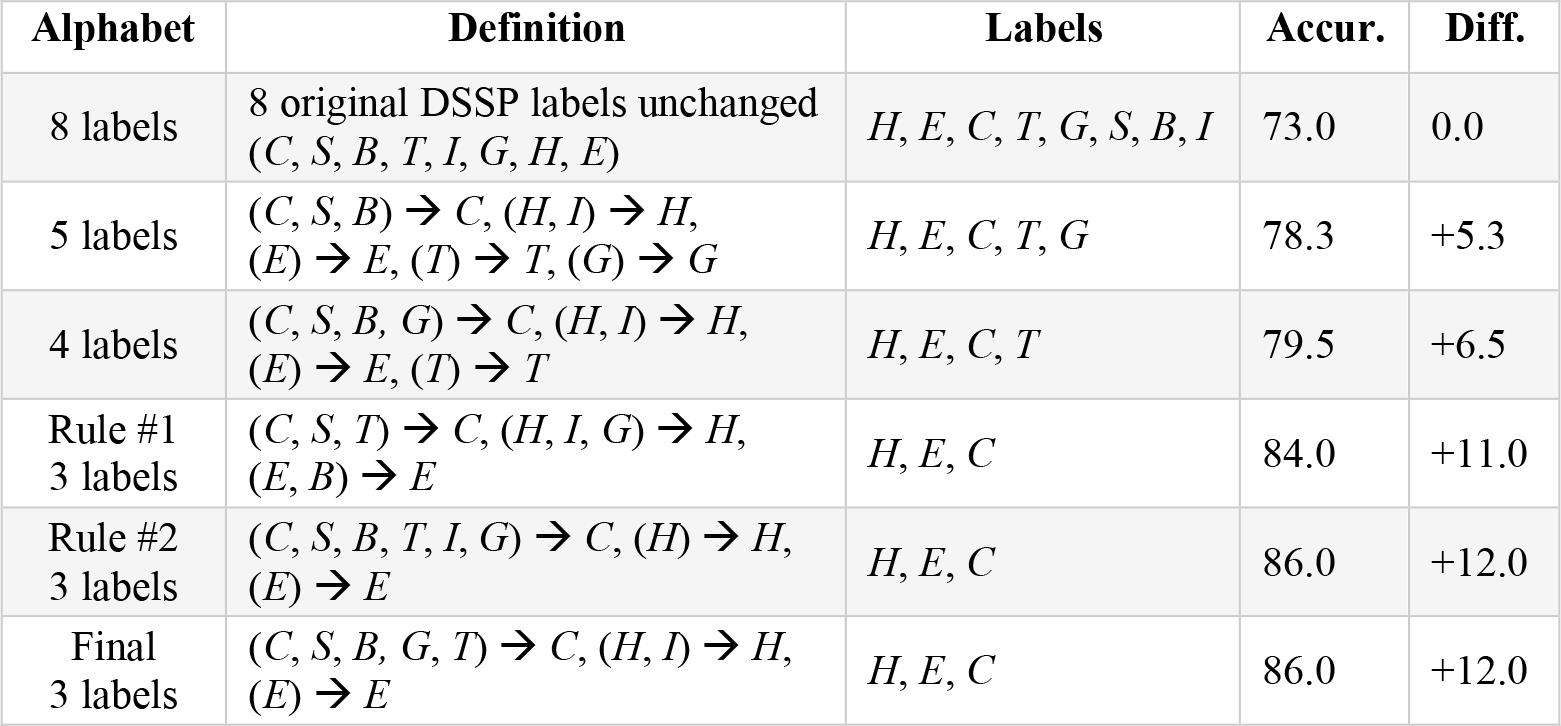
Label definitions for 3 classical and 3 new alphabets and associated SecNet accuracies on Test2018 **test set.**

| Alphabet | Definition | Labels | Accur. | Diff. |
| --- | --- | --- | --- | --- |
| 8 labels | 8 original DSSP labels unchanged<br>(*C, S, B, T, I, G, H, E*) | *H, E, C, T, G, S, B, I* | 73.0 | 0.0 |
| 5 labels | $(C, S, B) \rightarrow C, (H, I) \rightarrow H,$<br>$(E) \rightarrow E, (T) \rightarrow T, (G) \rightarrow G$ | *H, E, C, T, G* | 78.3 | +5.3 |
| 4 labels | $(C, S, B, G) \rightarrow C, (H, I) \rightarrow H,$<br>$(E) \rightarrow E, (T) \rightarrow T$ | *H, E, C, T* | 79.5 | +6.5 |
| Rule #1<br>3 labels | $(C, S, T) \rightarrow C, (H, I, G) \rightarrow H,$<br>$(E, B) \rightarrow E$ | *H, E, C* | 84.0 | +11.0 |
| Rule #2<br>3 labels | $(C, S, B, T, I, G) \rightarrow C, (H) \rightarrow H,$<br>$(E) \rightarrow E$ | *H, E, C* | 86.0 | +12.0 |
| Final<br>3 labels | $(C, S, B, G, T) \rightarrow C, (H, I) \rightarrow H,$<br>$(E) \rightarrow E$ | *H, E, C* | 86.0 | +12.0 |

#### Usage of our secondary-structure prediction software

Our secondary structure prediction software, *SecNet*, is free, open-source, and cross-platform. It is written in *Python* 3 and is downloadable as a single package from dunbrack.fccc.edu/ss and github.com/sh-maxim/ss with all required third-party software and databases included. When *SecNet* launches for the first time, it self-configures, detecting missing

*Python* libraries such as *numpy*, *keras*, *theano* or missing third-party databases, which it then automatically downloads and installs. The prediction mode of our application requires an average desktop with 2-4 CPU cores and 4-8 GB of RAM; during execution, it consumes ∼1-2 GB of RAM. For faster multi-threaded execution the software detects how many CPU cores are available and uses all of them unless fewer cores are assigned with an optional argument.

A user does not need to generate any input features manually; *SecNet* automatically prepares all required input features for a target protein sequence by running included third-party software with preprocessed sequence databases. For an average protein of 200-300 residues it takes about 10 minutes to prepare input features, several seconds to load predictive models from a storage device into memory, and a few seconds to run an ensemble of 10 NN models to perform secondary structure prediction. The usage is very simple with intuitive command-line arguments such as -label [3 or 4 or 5 or 8] and -rule1 or -rule2; the full instructions and options are available at the above online resources and also included in a readme file:

*secnet -help*

*secnet -i example.seq -label 8 -o example.ss8*

*secnet -input example.fasta -rule1 -label 3-output example.rule1ss3 secnet -i example.seq -label 4 -o example.ss4 -cpu 3*

## Discussion

We have developed *SecNet*, a simple traditional four-layer convolutional neural network for predicting protein secondary structure purely from sequence. We constructed new training, validation, and testing sets such that the test set consists of protein structures determined in 2018

January 1, 2018. This enabled fair and unbiased comparison of our method with two programs, *DeepCNF* and *eCRRNN*, which were trained on data released prior to 2018 and which had the highest reported accuracies on *CB513*, a widely used test set in secondary structure prediction studies. *SecNet* achieved accuracy of 73.0% on 8-label predictions, 84.0%, on *Rule #1* 3-label predictions, and 86% on *Rule #2* 3-label predictions on *Test2018*. It outperforms *DeepCNF* and *eCRRNN* by 2.0-3.4% on our *Test2018* set and *CB513*, and other methods by 1.7-8.8% on the *CB513* test set. While this is a small improvement in accuracy, it is significant in light of progress over the last 20 years—the accuracy of *PSIPRED*, published in 1999, on *CB513* was 79.2% for 3- label *Rule #1* predictions.

In the development of *SecNet*, we tried a variety of options for feature calculation, network structure, and training hyper-parameters. After selecting the best options with the validation set, we used the test set only once to estimate the accuracy. We then performed a retrospective ablation study, using the test set to calculate accuracy on the various networks produced during the earlier training and validation studies. The results not surprisingly indicate that the most accurate model is the result of many decisions each of which contributes toward the accuracy. The most interesting results of the ablation study were that window sizes larger than 29 residues did not increase the accuracy, and that *PSI-BLAST* profiles are more useful than HMMs in predicting secondary structure, even though the information in HMMs is inherently richer.

From our experience in developing the training, validation, and test set, we developed a short list of recommendations that might be considered in future efforts to avoid overtraining and to enable comparison of different programs:

### 1) Generating a list of chains for training, validation, and test sets

A majority of prediction methods enforce a maximal pairwise sequence similarity of 25% between training, validation, and/or test sets as well within the test set. But how the sequence identities are calculated can matter, and this is not always described in sufficient detail to ensure reproducibility. A few methods either do not list a threshold [77, 81] or use a higher value [82, 100], which may lead to a higher accuracy that would not be sustained on proteins unrelated to the training and testing sets [84]. For example, we observed that if our model is trained and tested on data sets with 50% sequence identity instead of 25%, it has 1.3% higher accuracy (top of Fig 4). This issue has been raised several times by different groups [34, 46, 90, 91]. In this work, we developed a rigorous protocol that relies on two passes of two different sequence alignment software programs (*HHblits* and *Clustal-Omega*) to enforce 25% identity within our data sets.

An unforeseen violation of a sequence-identity threshold may occur when a method relies on predictions from third-party software as input features when the third-party software was trained on protein structures that are more than 25% identical to sequences in the test set of the new method. This may occur for instance, if a method uses features such as predicted contacts or solvent exposure from other programs trained on sequences related to those in the test set. This can be avoided if the test set for the secondary structure prediction method contains sequences of structures determined that are less than 25% identical to the training sets of all predicted input features. This can be hard to determine, since not all training sets are defined by authors of feature prediction programs or there is a set of nested programs with each trained on its own training set. To avoid such a situation, we should enforce the pairwise similarity threshold between every new structure published after the third-party software release and those before it. It is necessary to compare all sequences before and after the required date, regardless of resolution, experimental method, or structure quality. These filters can be applied to the training, validation, and test data later.

Another consideration on which chains to include in the training, validation, and test sets is their secondary structure content. Alpha helices are easier to predict and a test set enriched with them is biased to have higher accuracy (Fig 3); therefore, data sets should have secondary structure labels representative of the general PDB population.

### 2) Sequences and labels for the training, validation, and test sets

Secondary structure cannot be assigned to amino acids with missing coordinates caused by unresolved electron density, and such amino acids are ignored by *DSSP*. A test set should consist of full sequences, and the labels therefore need to include a designation for residues missing from the coordinates. By contrast, the popular *CB513* test set does not include residues with missing coordinates. Information on disordered residues mapped to the sequence is provided in the *mmCIF* format of files from the PDB, or from the file “*ss_dis.txt*” available from the PDB (https://www.rcsb.org/pages/download/http#ss).

Previously secondary structure prediction methods are mostly trained to predict three labels: helix (*H*), sheet (*E*), and coil (*C*). In the early days, ground-truth labels came from secondary structure assigned by authors of deposited PDB structures. With emergence of a widely-adopted *DSSP* program in 1983, the 8 secondary-structure labels (*H*, *B*, *E*, *G*, *I*, *T*, *S, C*) assigned by this software have been converted into 3 labels (*H*, *E*, *C*); the sets of 8 and 3 labels are used separately for development of prediction methods. With a lack of standardization, methods report accuracies based on either *Rule #1*: (*H*, *G*, *I*) → *H*, (*E*, *B*) → *E*, (other 4 labels) → *C* or *Rule #2*: (*H*) → *H*, 1. (*E*) → *E*, (other 6 labels) → *C*. The 3-label *Rule #2* is easier for prediction than *Rule #1* by 2.0% (top of Fig 4). Some authors did not report which rule was used [73, 77, 78, 82]; some continue to use the easier *Rule #2* and compare their results to previous methods that used the harder *Rule #1* [81, 103]. Which rule is used should be clearly stated in a publication to avoid ambiguity.

Label sets greater in size than 3 but less than 8 may be useful in some contexts. We demonstrated that the *B* (beta bridge) and *S* (bend) labels have very little sequence signal, and since they may be difficult to sample anyway they can productively be converted to the generic *C* label. The *I* label (pi helices) is very rare, and may be converted to *H*, producing a 5-label rule: (*H*, *I*) → *H*; (*B*, *S*, *C*) → *C*; *E; T*; *G*. Residues in 310 helices are also difficult to predict. We showed that those 310 helix residues immediately adjacent to alpha helices have similar amino-acid preferences to alpha helices. Those isolated within coil regions, which are commonly of length 3, are mostly predicted as coil. However, these residues outnumber those adjacent to alpha helices, and *G* can therefore be productively converted to *C*. Therefore we define a 4-label rule as: (*H*, *I*) → *H*, (*B*, *S*, *C, G*) → *C; E; T*. Finally, we define a slightly modified version of *Rule #2*: (*H*, *I*) → *H*; (*B*, *S*, *C, T*) → *C; E*.

### 3) Separation of training, validation, and test sets

Some methods do not designate a separate validation set, and make design decisions based on repeated use of either the training set or a single test set. If a training set is used for these purposes, the produced model is over-trained and training accuracy will not propagate to the test set accuracy. For example, the accuracy on our validation set is 2.0% and 2.5% higher than that on the test sets for 3-label and 8-label predictions respectively. A validation set should be split from the training set and used for model selection, with the test set only used at the end to produce final accuracy estimations [104].

### 4) Distributing the data sets, including sequences and labels

The availability of reliable data sets is critical for reproducibility and fair benchmarking. For many publications, these sets are not readily available; contacting authors is time-consuming and often unproductive, and the needed information is not always still available. Data sets that can be reused by other groups to test new methods need to contain not only the PDB codes and chain identifiers but also the full sequences and label strings. The PDB can occasionally change a sequence and coordinates if an entry is updated. If the 8-label *DSSP* codes are converted to a smaller set, then providing the transformed labels is valuable to retain consistency. Complete data sets should be uploaded to at least two publicly available data sharing systems, for example to journal or institutional websites or *GitHub* or *DropBox* or *SourceForge*. A journal may not always provide adequate storage for bulky data sets.

## Methods

### Protocol for preparation of 2018 data sets

A complete list of PDB entries was obtained from PISCES [27] server on November 12, 2018. The entry list was divided into *Before18*, which consists of entries released prior to January 1, 2018, and *After18* which consists of those released on or after the same date (Fig 1). Chain sequences were obtained from PISCES. Identical sequences were removed from *Before18* and *After18*, and sequences identical to any sequence in *Before18* were removed from *After18*. Each identical sequence was linked with a complete list of its protein structures of varying quality.

We used the output of two sequence-alignment programs, *HHblits* and *Clustal-Omega*, to calculate pairwise sequence similarities within and between the sequences in the *Before18* and

*After18* sets. The *HHblits* pairwise sequence identities were calculated for use in the PISCES server. Any item in *After18* was removed if it had more than 25% sequence identity to any item in *Before18* calculated with either *HHblits* or *Clustal-Omega*. We underline that *Before18* includes PDB sequences of any experiment or resolution to eliminate any test sequence too similar even to a low-quality pre-2018 sequence that could have been included in training of previous methods. Usage of two programs helps if either one fails to detect similarity.

We applied several filters to the structures and sequences in *Before2018* and *After2018* before applying a pairwise sequence identity filter within each set. We removed structures having resolution worse than 2.2 Å, or free R-factor greater than 0.25 (if free-R factor was not available, we used the R-factor+0.05) [92]. We excluded chains of length less than 40 residues and Cα-atom- only models. Next we enforced 25% pairwise identity in each set based on the sequence identities obtained from *HHblits* and *Clustal-Omega*. We removed any sequences from *Before2018* that was more than 25% identical with any sequence in *CB513* so that we could use *Set2018* to train *SecNet* and use *CB513* as an additional test set (*Test2018* already cannot have sequences more than 25% identical to the structures in *CB513* that were determined before the year 2000). For identical sequences, we selected a protein chain from a crystal structure having the highest resolution, followed by the best R-factor, and then the PDB chain having the largest number of coordinates.

This procedure produced the *Test2018* test set (*After2018* after the quality and sequence identity filters were applied), which contains sequences that are not more than 25% similar to *any* chain in a PDB entry released before Jan 1, 2018. Similarly, the *Set2018* training and validation sets were created from a 9 to 1 random split of the chains in *Before2018* after the quality and sequence identity filters were applied. We note that complete chain sequences are stored (not just the ones with coordinates) in our sets. We store original one-letter sequences using the standard representation where a non-standard amino acid has a single-letter code of the closest analog among 20 standard amino acids if it exists; otherwise “X” is used.

#### Input labels and features

The 8 secondary-structure labels were obtained from the file, “*ss_dis.txt*” which has records for the entire PDB and is available for download from PDB. This file also contains strings that indicate the ordered and disordered residues in the sequence. The *DSSP* and disorder strings were combined, with spaces in the *DSSP* string replaced with “C” when the disorder string indicated an ordered residue (“-“) and “X” when the disorder string indicated a disordered residue (“X”). For instance, for PDB entry 1A64 chain A, this file contains:

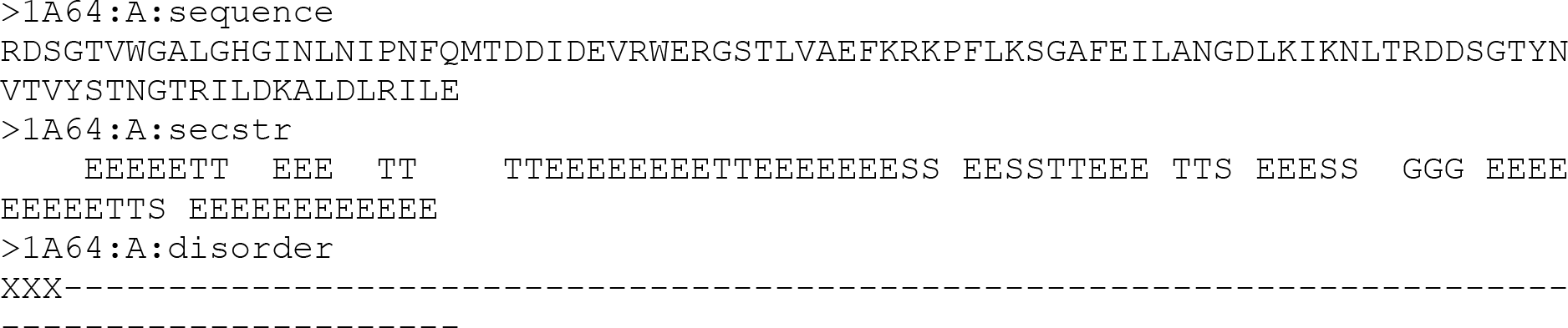

which results in the combined *DSSP*-disorder string:

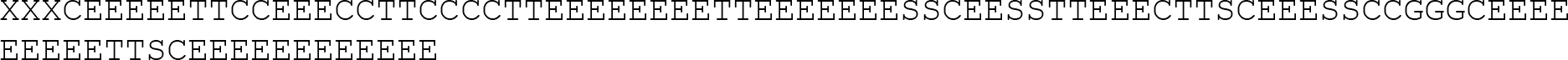

Training was performed and test accuracy was reported on all residues with known coordinates for which *DSSP* was able to assign labels, i.e. for residues with ordered ‘*-*’ flag and not with disorder ‘*X*’ flag. Each amino acid subject to training or testing includes a full amino-acid sequence around it including disordered residues.

In addition to the labels, the input to our CNN includes a 92-by-29 feature matrix where the columns represent 14 residues to the left from the central residue, the central residue which is subject to secondary structure prediction, and 14 residues to the right; the rows consist of 92 features for each position of 29 residues. Among the features are 1) one-hot encoding of 22 residue types: the 20 standard amino acids, “X” (a non-standard residue without a standard amino-acid analog), and “?” which denotes a non-existing residue outside the sequence when the central residue of an input window is too close to the beginning and end of the sequence; 2) two sets of *PsiBlast* sequence profiles generated against *Uniref90* sequence database at the end of the first and second rounds, each 20 long – the number of standard amino acids; 3) 30 parameters of Hidden Markov Model of the target generated by a search against the *uniprot20* sequence database computed with *hhsuite*. Thus, the number of rows in the input matrix is 22 + 2 * 20 + 30 = 92.

*PsiBlast* version 2.6.0 was obtained from NCBI, and used to search the *Uniref90* database of 58,284,765 sequences downloaded on Aug 2, 2017 from www.uniprot.org/downloads. *Uniref90* was converted to a *PsiBlast* database with the command: makeblastdb -in uniref90.fasta -parse_seqids -dbtype prot The *PsiBlast searches were performed with the command: siblast -db uniref90.binary -query example.fasta-inclusion_ethresh 0.001-evalue 10-save_pssm_after_last_round-out_ascii_pssm example.mtx-num_iterations 2-num_threads 16-save_each_pssm*

The last flag forces *PsiBlast* to output each PSSM with a different file name (−1, −2, etc.). The input to *SecNet* consists of the output log-scale scores instead of rounded weighted percentages. For non-existing residues outside the complete sequence, log score values of −19 were used at each of the 20 standard amino-acid positions.

The HMM of a target vs. *uniprot20* sequence database was created with 2.0.16 *hhsuite*. *uniprot20* is located at www.user.gwdg.de/~compbiol/data/hhsuite/databases/hhsuite_dbs/old- releases and dated Feb, 2016. All 30 parameters of each residue in a target sequence are used from the HMM file generated with a set of two commands:

### hhblits −i example.fasta -d uniprot20 -oa3m example.a3m -cpu 16 hhmake -i example.a3m -o example.hhm

The 30 parameters with ‘*\**’ values representing +∞ values were replaced with 99999.0. For the non-existing residues outside the *N* and *C* termini, the first 27 variables (*A*’, ‘*C*’, ‘*D*’, ‘*E*’, ‘*F*’, ‘*G*’, ‘*H*’, ‘*I*’, ‘*K*’, ‘*L*’, ‘*M*’, ‘*N*’, ‘*P*’, ‘*Q*’, ‘*R*’, ‘*S*’, ‘*T*’, ‘*V*’, ‘*W*’, ‘*Y*’, ‘*M*→*M*’, ‘*M*→*I*’, ‘*M*→*D*’, ‘*I*→*M*’, ‘*I*→*I*’, ‘*D*→*M*’, ‘*D*→*D*’) of 99999.0 value and the last 3 variables (‘*Neff*’, ‘*Neff_I*’, ‘*Neff_D*’) of 0.0 value were used for HMM parameterization. At the end all HMM parameters were divided by a normalization factor of 99,999.0.

### Training and testing a CNN for secondary structure prediction

Our neural network is a traditional CNN (Fig 2) consisting of an input layer which reads a 92-by-29 matrix centered on a residue subject to secondary structure label prediction, 4 convolutional hidden layers, densely-connected NN layer with an output dimensionality of the number of secondary structure labels, and a *softmax* activation layer returning a vector of probabilities for each label. The total number of *CNN* parameters is 2,397,960. The 4 convolutional hidden layers have the same filter length of 7 with valid border treatment and a rectifier activation function; the number of filters at each hidden layer increases by 128 starting from 128 at the first hidden layer: 128, 256, 384 and 512. As a training regularization technique, a dropout layer with 0.30 value is implemented in front of each of the 4 hidden layers and the dense layer. This CNN was implemented and trained using 1.2.2 *Keras*, an open-source NN *Python* library with 0.8.2 *Theano* backend.

We independently trained 10 CNN models by 10-fold cross-validation on the training set. To gain additional accuracy improvement, we used a popular technique where a final predictive model is an ensemble of 10 CNNs returning a majority label vote based on 10 outputs of a predicted label. The final accuracy was reported based the independent test set that had never been used during training, validation, or making design decisions. The test set was used only once to report the final results such as the final overall accuracy, confusion matrix, and accuracy improvement for each component of the predictive model during the retrospective ablation study to update these overestimated values based on the original validation set with the unbiased ones based on the testing set. However, during algorithm development all decisions on how to train, what input features to use, what third-party software arguments to use, which sequence databases to employ and what CNN architecture to use were only based on the validation set to avoid overestimated accuracy at the very end with the testing set.

A stochastic gradient descent was used for optimization. We tried many different combinations of parameters and empirically selected the following optimization parameters leading to the best validation accuracy during many training trials: *loss function = categorical cross entropy*, *number of epochs = 650*, *batch size = 256*, *learning rate = 0.01*, *momentum = 0.9*, *Nesterov accelerated gradient = off*, *decay = learning rate / number of epochs = 0.01 / 650 = 1.54e-5* and *dropout = 0.3*. We computed validation accuracy at the end of each epoch and saved a model if the validation accuracy was higher than at any previous epoch disregarding a value of the training set accuracy.

A single training requires about 4 days on a 16-core CPU workstation; within the first 12- 24 hours the validation accuracy was typically 0.3-0.5% below the final best accuracy. The training was stable if different starting random seed states were used leading to a spread of final validation accuracies of only up to 0.1%.

Training and testing were based on overall accuracy which includes all residues with assigned secondary structure different from ‘*X*’ (disordered residues with missing coordinates). The accuracy was calculated as a number of correctly predicted labels / number of all assigned labels.

### Benchmarking of competitor software

We ran *DeepCNF* and *eCRRNN* with default arguments. An ensemble of 10 models was enabled for *eCRRNN*. We made sure that we were executing these third-party programs properly by benchmarking them on data sets included in their respective published studies and reproducing the same results within 0.1% accuracy for each included set. As with *SecNet*, *DeepCNF* and *eCRRNN* were benchmarked on complete original sequences and accuracy was reported on residues of known secondary structure.

## Supporting information

S2 Data set. Tab-delimited Set2018 (PDB, chain ID, sequence, DSSP sec. struc. and converted to 3, 4 & 5 labels) for train, valid, test (Test2018) sets

## Supporting information

**S1 Supporting information**. This file contains supplementary Figure A, Table A and Table B.

**S2 Data set**. This tab-delimited plain text file contains *Set 2018* data set which includes PDB entry codes, chain IDs, complete amino-acid sequences and ground-truth secondary structure information (*DSSP* 8 labels, *Rule #1* and *Rule #2* 3 labels, 4 labels, and 5 labels) for the training, validation, and testing (*Test2018*) sets.

---

**Fig A.**
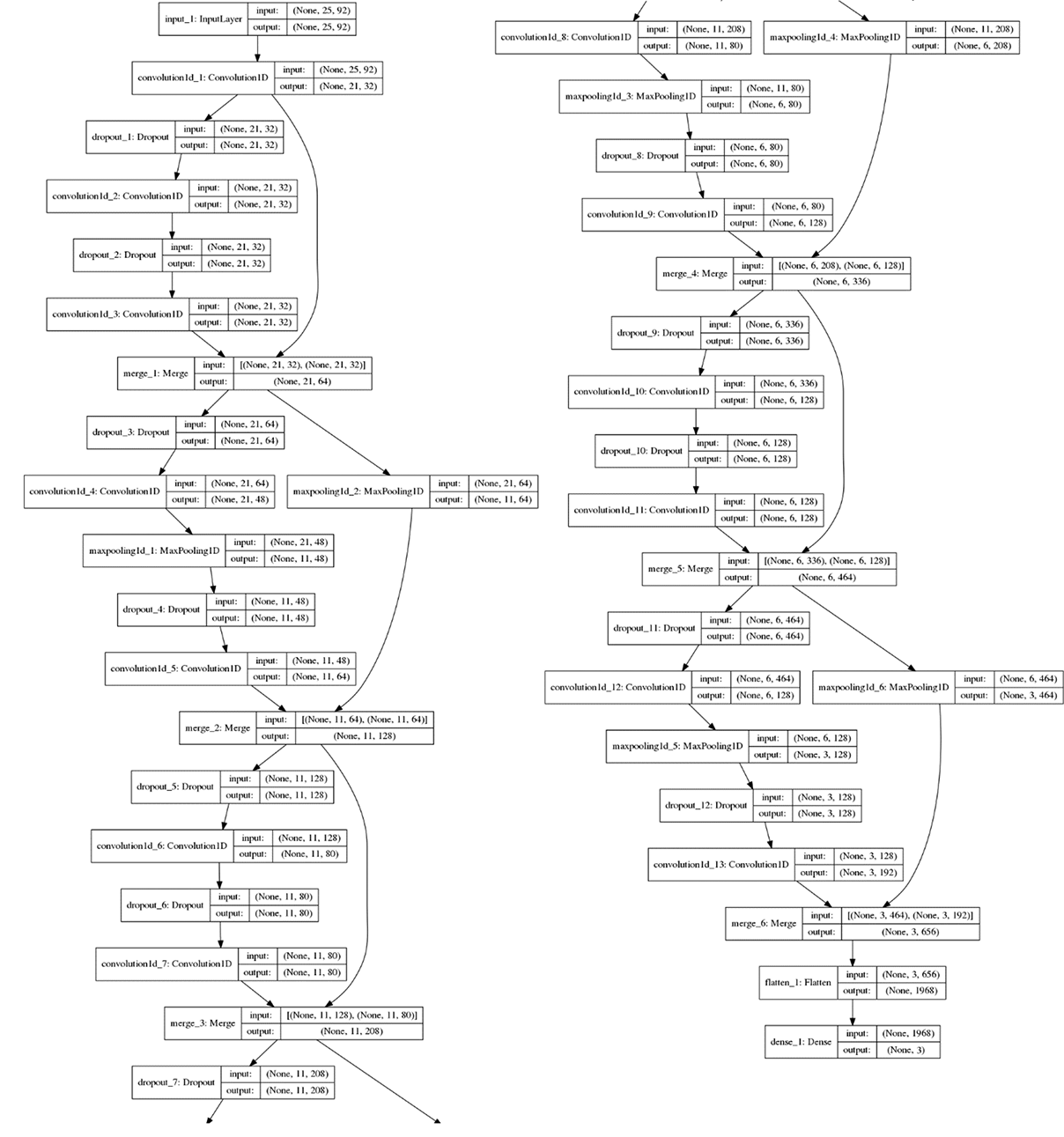
Architecture of *SecRes*, our abandoned Residual Neural Network with input window of 21 to 51 amino acids and 20-40 convolutional layers. A sample architecture diagram of our residual neural network, *SecRes* from a family of networks with input of 21 to 51 amino acids and 20 to 40 layers. The sample has 13 hidden convolutional layers, 6 shortcut connections each bypassing 2 hidden layers and input window of 25 amino acids. Output from the blocks of 2 bypassed hidden layers is concatenated with the input to these blocks. Maxpooling of size 2 is applied for linear dimensionality reduction. Other details about the network are same as in Fig 2 about our traditional 4-layer CNN *SecNet*. We varied network complexity with a layer number, input size, and number of training parameters from 600 thousand to 20 million, observed same or worse accuracy as for *SecNet*, and as a result abandoned this more complex network.

**Table A.**
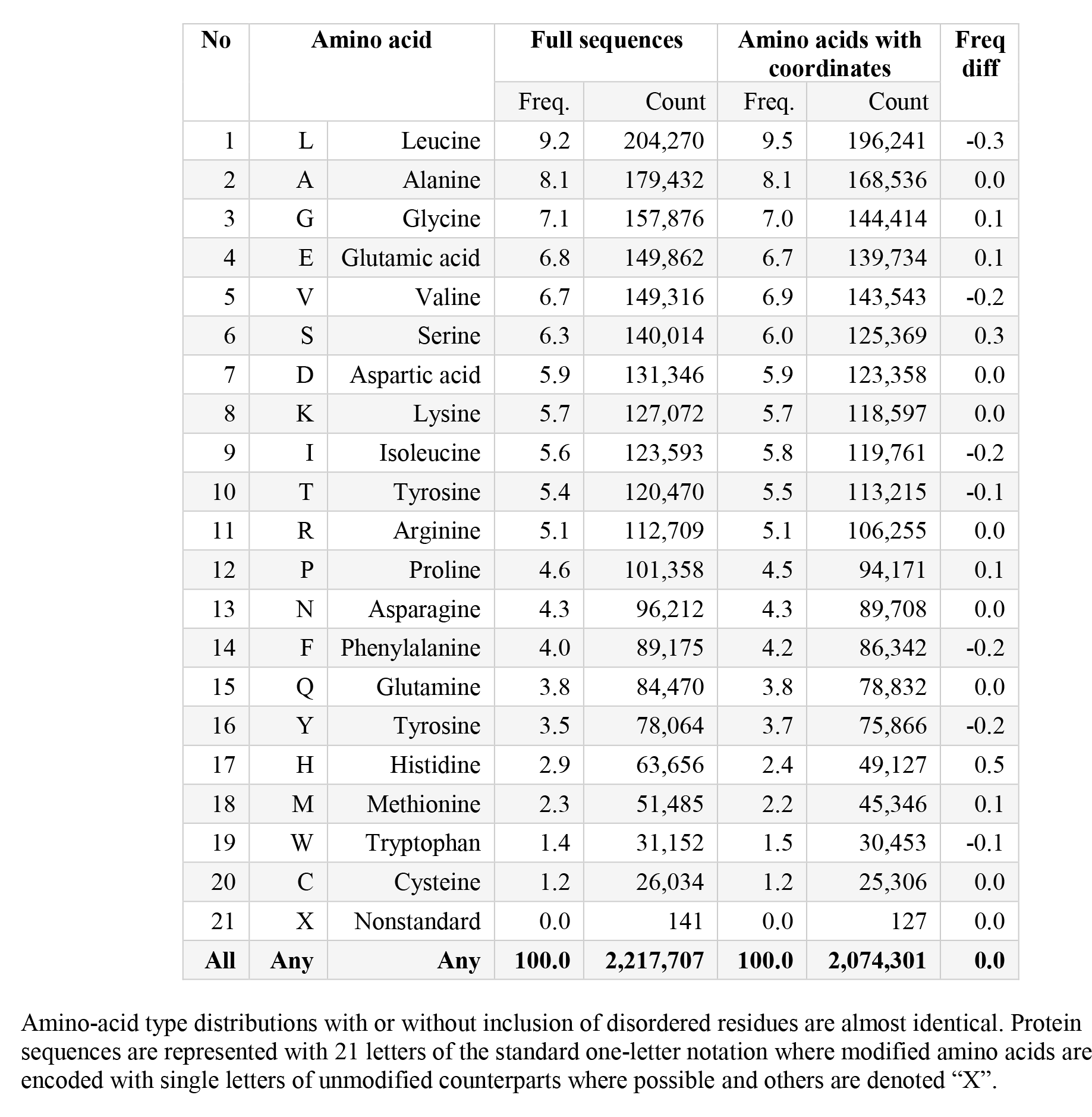
Amino-acid frequencies in 8,712 proteins of Set2018.

**Table B.**
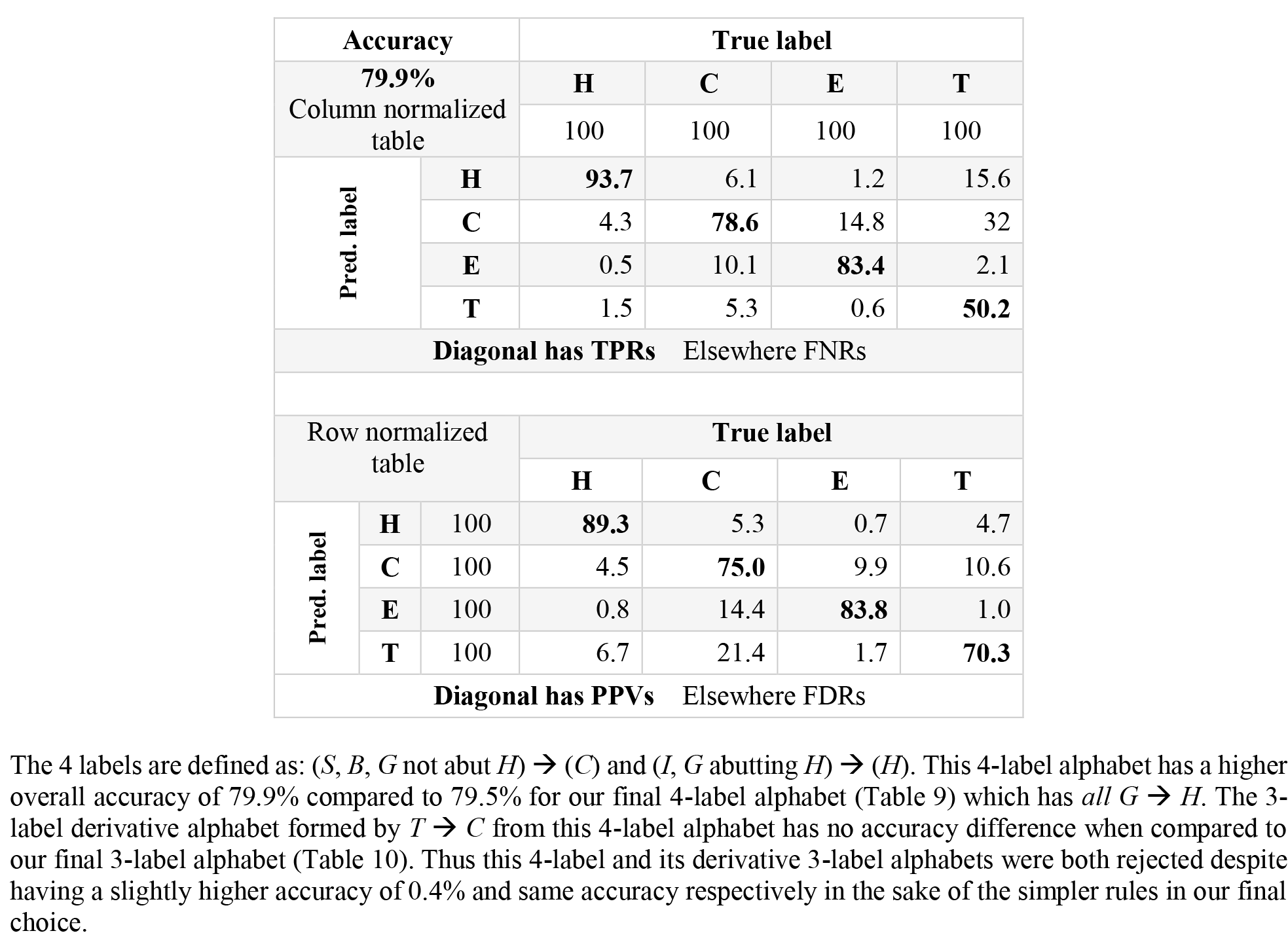
Alternative 4-label alphabet: H, C, E, and T where G abutting H → H and rest G → C: (1) recalls and false negative rates and (2) precisions and false discover rates of SecNet.

## References

1. Yang Y, Gao J, Wang J, Heffernan R, Hanson J, Paliwal K, et al. Sixty-five years of the long march in protein secondary structure prediction: the final stretch? Brief Bioinform. 2018;19(3):482–94. Epub 2017/01/04. doi: 10.1093/bib/bbw129. PubMed PMID: 28040746; PubMed Central PMCID: PMCPMC5952956.

2. Kabsch W, Sander C. Dictionary of protein secondary structure: pattern recognition of hydrogen-bonded and geometrical features. Biopolymers. 1983;22(12):2577–637. Epub 1983/12/01. doi: 10.1002/bip.360221211. PubMed PMID: 6667333.

3. Fischer D, Eisenberg D. Protein fold recognition using sequence-derived predictions. Protein Sci. 1996;5(5):947–55. Epub 1996/05/01. doi: 10.1002/pro.5560050516. PubMed PMID: 8732766; PubMed Central PMCID: PMCPMC2143416.

4. Skolnick J, Kihara D, Zhang Y. Development and large scale benchmark testing of the PROSPECTOR_3 threading algorithm. Proteins. 2004;56(3):502–18. Epub 2004/07/02. doi: 10.1002/prot.20106. PubMed PMID: 15229883.

5. Rohl CA, Strauss CE, Misura KM, Baker D. Protein structure prediction using Rosetta. Methods Enzymol. 2004;383:66–93. Epub 2004/04/06. doi: 10.1016/S0076-6879(04)83004-0. PubMed PMID: 15063647.

6. Wu S, Skolnick J, Zhang Y. Ab initio modeling of small proteins by iterative TASSER simulations. BMC Biol. 2007;5:17. Epub 2007/05/10. doi: 10.1186/1741-7007-5-17. PubMed PMID: 17488521; PubMed Central PMCID: PMCPMC1878469.

7. Kamisetty H, Ovchinnikov S, Baker D. Assessing the utility of coevolution-based residue-residue contact predictions in a sequence- and structure-rich era. Proc Natl Acad Sci U S A. 2013;110(39):15674–9. Epub 2013/09/07. doi: 10.1073/pnas.1314045110. PubMed PMID: 24009338; PubMed Central PMCID: PMCPMC3785744.

8. Adhikari B, Bhattacharya D, Cao R, Cheng J. CONFOLD: Residue-residue contact- guided ab initio protein folding. Proteins. 2015;83(8):1436–49. Epub 2015/05/15. doi: 10.1002/prot.24829. PubMed PMID: 25974172; PubMed Central PMCID: PMCPMC4509844.

9. Ovchinnikov S, Park H, Varghese N, Huang PS, Pavlopoulos GA, Kim DE, et al. Protein structure determination using metagenome sequence data. Science. 2017;355(6322):294-8. Epub 2017/01/21. doi: 10.1126/science.aah4043. PubMed PMID: 28104891; PubMed Central PMCID: PMCPMC5493203.

10. Plaxco KW, Simons KT, Baker D. Contact order, transition state placement and the refolding rates of single domain proteins. J Mol Biol. 1998;277(4):985–94. Epub 1998/05/30. doi: 10.1006/jmbi.1998.1645. PubMed PMID: 9545386.

11. Ahmad S, Gromiha MM, Sarai A. Real value prediction of solvent accessibility from amino acid sequence. Proteins. 2003;50(4):629–35. Epub 2003/02/11. doi: 10.1002/prot.10328. PubMed PMID: 12577269.

12. Adamczak R, Porollo A, Meller J. Accurate prediction of solvent accessibility using neural networks-based regression. Proteins. 2004;56(4):753–67. Epub 2004/07/29. doi: 10.1002/prot.20176. PubMed PMID: 15281128.

13. Heffernan R, Dehzangi A, Lyons J, Paliwal K, Sharma A, Wang J, et al. Highly accurate sequence-based prediction of half-sphere exposures of amino acid residues in proteins. Bioinformatics. 2016;32(6):843–9. Epub 2015/11/17. doi: 10.1093/bioinformatics/btv665. PubMed PMID: 26568622.

14. Kaur H, Raghava GP. A neural network method for prediction of beta-turn types in proteins using evolutionary information. Bioinformatics. 2004;20(16):2751–8. Epub 2004/05/18. doi: 10.1093/bioinformatics/bth322. PubMed PMID: 15145798.

15. Petersen B, Lundegaard C, Petersen TN. NetTurnP--neural network prediction of beta-turns by use of evolutionary information and predicted protein sequence features. PLoS One. 2010;5(11):e15079. Epub 2010/12/15. doi: 10.1371/journal.pone.0015079. PubMed PMID: 21152409; PubMed Central PMCID: PMCPMC2994801.

16. Kountouris P, Hirst JD. Predicting beta-turns and their types using predicted backbone dihedral angles and secondary structures. BMC Bioinformatics. 2010;11:407. Epub 2010/08/03. doi: 10.1186/1471-2105-11-407. PubMed PMID: 20673368; PubMed Central PMCID: PMCPMC2920885.

17. Schlessinger A, Rost B. Protein flexibility and rigidity predicted from sequence. Proteins. 2005;61(1):115–26. Epub 2005/08/05. doi: 10.1002/prot.20587. PubMed PMID: 16080156.

18. Radivojac P, Iakoucheva LM, Oldfield CJ, Obradovic Z, Uversky VN, Dunker AK. Intrinsic disorder and functional proteomics. Biophys J. 2007;92(5):1439-56. Epub 2006/12/13. doi: 10.1529/biophysj.106.094045. PubMed PMID: 17158572; PubMed Central PMCID: PMCPMC1796814.

19. Simossis VA, Heringa J. Integrating protein secondary structure prediction and multiple sequence alignment. Curr Protein Pept Sci. 2004;5(4):249–66. Epub 2004/08/24. PubMed PMID: 15320732.

20. Zhou H, Zhou Y. SPEM: improving multiple sequence alignment with sequence profiles and predicted secondary structures. Bioinformatics. 2005;21(18):3615–21. Epub 2005/07/16. doi: 10.1093/bioinformatics/bti582. PubMed PMID: 16020471.

21. Zhang T, Faraggi E, Li Z, Zhou Y. Intrinsically semi-disordered state and its role in induced folding and protein aggregation. Cell Biochem Biophys. 2013;67(3):1193–205. Epub 2013/06/01. doi: 10.1007/s12013-013-9638-0. PubMed PMID: 23723000; PubMed Central PMCID: PMCPMC3838602.

22. Pei J, Kim BH, Grishin NV. PROMALS3D: a tool for multiple protein sequence and structure alignments. Nucleic Acids Res. 2008;36(7):2295–300. Epub 2008/02/22. doi: 10.1093/nar/gkn072. PubMed PMID: 18287115; PubMed Central PMCID: PMCPMC2367709.

23. Godzik A, Jambon M, Friedberg I. Computational protein function prediction: are we making progress? Cell Mol Life Sci. 2007;64(19-20):2505–11. Epub 2007/07/06. doi: 10.1007/s00018-007-7211-y. PubMed PMID: 17611711.

24. Taherzadeh G, Zhou Y, Liew AW, Yang Y. Sequence-Based Prediction of Protein-Carbohydrate Binding Sites Using Support Vector Machines. J Chem Inf Model. 2016;56(10):2115–22. Epub 2016/10/25. doi: 10.1021/acs.jcim.6b00320. PubMed PMID: 27623166.

25. Li B, Krishnan VG, Mort ME, Xin F, Kamati KK, Cooper DN, et al. Automated inference of molecular mechanisms of disease from amino acid substitutions. Bioinformatics. 2009;25(21):2744–50. Epub 2009/09/08. doi: 10.1093/bioinformatics/btp528. PubMed PMID: 19734154; PubMed Central PMCID: PMCPMC3140805.

26. Guzzo AV. The influence of amino acid sequence on protein structure. Biophysical journal. 1965;5(6):809–22.

27. Wang G, Dunbrack RL, Jr. PISCES: a protein sequence culling server. Bioinformatics. 2003;19(12):1589–91. Epub 2003/08/13. doi: 10.1093/bioinformatics/btg224. PubMed PMID: 12912846.

28. Jones DT. Protein secondary structure prediction based on position-specific scoring matrices. J Mol Biol. 1999;292(2):195–202.

29. Heringa J. Computational methods for protein secondary structure prediction using multiple sequence alignments. Curr Protein Pept Sci. 2000;1(3):273–301. Epub 2002/10/09. PubMed PMID: 12369910.

30. Rost B. Review: protein secondary structure prediction continues to rise. J Struct Biol. 2001;134(2-3):204–18. Epub 2001/09/12. doi: 10.1006/jsbi.2001.4336. PubMed PMID: 11551180.

31. Yoo PD, Zhou BB, Zomaya AY. Machine learning techniques for protein secondary structure prediction: An overview and evaluation. Curr Bioinform. 2008;3(2):74–86. doi: Doi 10.2174/157489308784340676. PubMed PMID: WOS:000255682100002.

32. Pirovano W, Heringa J. Protein secondary structure prediction. Methods Mol Biol. 2010;609:327–48. Epub 2010/03/12. doi: 10.1007/978-1-60327-241-4_19. PubMed PMID: 20221928.

33. Zhou Y, Faraggi E. In: Rangwala H, Karypis G, editors. Protein Structure Methods and Algorithms. Hoboken: John Wiley & Sons; 2010. p. 44–74.

34. Rost B, Sander C. Third Generation Prediction of Secondary Structures. In: Webster DM, editor. Protein Structure Prediction: Methods and Protocols. Totowa, NJ: Humana Press; 2000. p. 71-95.

35. Chou PY, Fasman GD. Prediction of protein conformation. Biochemistry. 1974;13(2):222–45. Epub 1974/01/15. PubMed PMID: 4358940.

36. Lim VI. Structural principles of the globular organization of protein chains. A stereochemical theory of globular protein secondary structure. J Mol Biol. 1974;88(4):857–72. Epub 1974/10/05. PubMed PMID: 4427383.

37. Garnier J, Osguthorpe DJ, Robson B. Analysis of the accuracy and implications of simple methods for predicting the secondary structure of globular proteins. J Mol Biol. 1978;120(1):97–120. Epub 1978/03/25. PubMed PMID: 642007.

38. Kabsch W, Sander C. How Good Are Predictions of Protein Secondary Structure. Febs Lett. 1983;155(2):179–82. doi: Doi 10.1016/0014-5793(82)80597-8. PubMed PMID: WOS:A1983QS70100001.

39. Dickerson RE, Timkovich R, Almassy RJ. The cytochrome fold and the evolution of bacterial energy metabolism. J Mol Biol. 1976;100(4):473–91. Epub 1976/02/05. PubMed PMID: 176369.

40. Schneider R. Sekundärstrukturvorhersage von Proteinen unter Berücksichtigung von Tertiärstrukturaspekten.: Diploma thesis: Department of Biology, University of Heidelberg, Heidelberg, Germany; 1989.

41. Ptitsyn OB, Finkelstein AV. Theory of protein secondary structure and algorithm of its prediction. Biopolymers. 1983;22(1):15–25. Epub 1983/01/01. doi: 10.1002/bip.360220105. PubMed PMID: 6673754.

42. Garnier J, Gibrat JF, Robson B. GOR method for predicting protein secondary structure from amino acid sequence. Methods Enzymol. 1996;266:540–53. Epub 1996/01/01. PubMed PMID: 8743705.

43. Gibrat JF, Garnier J, Robson B. Further developments of protein secondary structure prediction using information theory. New parameters and consideration of residue pairs. J Mol Biol. 1987;198(3):425–43. Epub 1987/12/05. PubMed PMID: 3430614.

44. Kabsch WS, C. Segment83 (unpublished). 1983.

45. Zvelebil MJ, Barton GJ, Taylor WR, Sternberg MJ. Prediction of protein secondary structure and active sites using the alignment of homologous sequences. J Mol Biol. 1987;195(4):957–61. Epub 1987/06/20. PubMed PMID: 3656439.

46. Rost B. PHD: predicting one-dimensional protein structure by profile-based neural networks. Methods Enzymol. 1996;266:525–39. Epub 1996/01/01. PubMed PMID: 8743704.

47. Levin JM, Pascarella S, Argos P, Garnier J. Quantification of secondary structure prediction improvement using multiple alignments. Protein Eng. 1993;6(8):849–54. Epub 1993/11/01. PubMed PMID: 8309932.

48. Solovyev VV, Salamov AA. Predicting alpha-helix and beta-strand segments of globular proteins. Comput Appl Biosci. 1994;10(6):661–9. Epub 1994/12/01. PubMed PMID: 7704665.

49. Jones DT. Protein secondary structure prediction based on position-specific scoring matrices. J Mol Biol. 1999;292(2):195–202. Epub 1999/09/24. doi: 10.1006/jmbi.1999.3091. PubMed PMID: 10493868.

50. Cuff JA, Clamp ME, Siddiqui AS, Finlay M, Barton GJ. JPred: a consensus secondary structure prediction server. Bioinformatics. 1998;14(10):892–3. Epub 1999/02/03. doi: 10.1093/bioinformatics/14.10.892. PubMed PMID: 9927721.

51. Pollastri G, Przybylski D, Rost B, Baldi P. Improving the prediction of protein secondary structure in three and eight classes using recurrent neural networks and profiles. Proteins. 2002;47(2):228–35. Epub 2002/04/05. PubMed PMID: 11933069.

52. Rost B, Sander C. Combining evolutionary information and neural networks to predict protein secondary structure. Proteins. 1994;19(1):55–72. Epub 1994/05/01. doi: 10.1002/prot.340190108. PubMed PMID: 8066087.

53. Cuff JA, Barton GJ. Application of multiple sequence alignment profiles to improve protein secondary structure prediction. Proteins. 2000;40(3):502–11. Epub 2000/06/22. PubMed PMID: 10861942.

54. Baldi P, Brunak S, Frasconi P, Soda G, Pollastri G. Exploiting the past and the future in protein secondary structure prediction. Bioinformatics. 1999;15(11):937–46. Epub 2000/04/01. PubMed PMID: 10743560.

55. Figureau A, Soto MA, Toha J. A pentapeptide-based method for protein secondary structure prediction. Protein Eng. 2003;16(2):103–7. Epub 2003/04/05. doi: 10.1093/proeng/gzg019. PubMed PMID: 12676978.

56. Kilosanidze GT, Kutsenko AS, Esipova NG, Tumanyan VG. Analysis of forces that determine helix formation in alpha-proteins. Protein Sci. 2004;13(2):351–7. Epub 2004/01/24. doi: 10.1110/ps.03429104. PubMed PMID: 14739321; PubMed Central PMCID: PMCPMC2286714.

57. Woo SK, Park CB, Lee SW. Protein secondary structure prediction using sequence profile and conserved domain profile. Lect Notes Comput Sc. 2005;3645:1–10. PubMed PMID: WOS:000232529000001.

58. Birzele F, Kramer S. A new representation for protein secondary structure prediction based on frequent patterns. Bioinformatics. 2006;22(21):2628–34. doi: 10.1093/bioinformatics/btl453. PubMed PMID: WOS:000241629600008.

59. Mooney C, Vullo A, Pollastri G. Protein structural motif prediction in multidimensional phi-psi space leads to improved secondary structure prediction. J Comput Biol. 2006;13(8):1489–502. doi: Doi 10.1089/cmb.2006.13.1489. PubMed PMID: WOS:000241815900007.

60. Wood MJ, Hirst JD. Protein secondary structure prediction with dihedral angles. Proteins-Structure Function and Bioinformatics. 2005;59(3):476–81. doi: 10.1002/prot.20435. PubMed PMID: WOS:000228779200007.

61. Midic U, Dunker AK, Obradovic Z. Exploring alternative knowledge representations for protein secondary-structure prediction. Int J Data Min Bioin. 2007;1(3):286–313. doi: Doi 10.1504/Ijdmb.2007.011614. PubMed PMID: WOS:000247735600005.

62. Momen-Roknabadi A, Sadeghi M, Pezeshk H, Marashi SA. Impact of residue accessible surface area on the prediction of protein secondary structures. Bmc Bioinformatics. 2008;9. doi: Artn 357 10.1186/1471-2105-9-357. PubMed PMID: WOS:000259530900001.

63. Meiler J, Baker D. Coupled prediction of protein secondary and tertiary structure. P Natl Acad Sci USA. 2003;100(21):12105–10. doi: 10.1073/pnas.1831973100. PubMed PMID: WOS:000186024300034.

64. Gassend B, O’Donnell CW, Thies W, Lee A, van Dijk M, Devadas S. Learning biophysically-motivated parameters for alpha helix prediction. Bmc Bioinformatics. 2007;8. doi: Artn S3 10.1186/1471-2105-8-S5-S3. PubMed PMID: WOS:000247560800003.

65. Meiler J, Muller M, Zeidler A, Schmaschke F. Generation and evaluation of dimension- reduced amino acid parameter representations by artificial neural networks. J Mol Model. 2001;7(9):360–9. doi: Doi 10.1007/s008940100038. PubMed PMID: WOS:000171594100006.

66. Adamczak R, Porollo A, Meller J. Combining prediction of secondary structure and solvent accessibility in proteins. Proteins. 2005;59(3):467–75. Epub 2005/03/16. doi: 10.1002/prot.20441. PubMed PMID: 15768403.

67. Wood MJ, Hirst JD. Protein secondary structure prediction with dihedral angles. Proteins. 2005;59(3):476–81. Epub 2005/03/22. doi: 10.1002/prot.20435. PubMed PMID: 15778963.

68. Dor O, Zhou Y. Achieving 80% ten-fold cross-validated accuracy for secondary structure prediction by large-scale training. Proteins. 2007;66(4):838–45. Epub 2006/12/21. doi: 10.1002/prot.21298. PubMed PMID: 17177203.

69. Lin HN, Chang JM, Wu KP, Sung TY, Hsu WL. HYPROSP II--a knowledge-based hybrid method for protein secondary structure prediction based on local prediction confidence. Bioinformatics. 2005;21(15):3227–33. Epub 2005/06/04. doi: 10.1093/bioinformatics/bti524. PubMed PMID: 15932901.

70. Montgomerie S, Sundararaj S, Gallin WJ, Wishart DS. Improving the accuracy of protein secondary structure prediction using structural alignment. BMC Bioinformatics. 2006;7:301. Epub 2006/06/16. doi: 10.1186/1471-2105-7-301. PubMed PMID: 16774686; PubMed Central PMCID: PMCPMC1550433.

71. Bondugula R, Xu D. MUPRED: a tool for bridging the gap between template based methods and sequence profile based methods for protein secondary structure prediction. Proteins. 2007;66(3):664–70. Epub 2006/11/17. doi: 10.1002/prot.21177. PubMed PMID: 17109407.

72. Pollastri G, Martin AJ, Mooney C, Vullo A. Accurate prediction of protein secondary structure and solvent accessibility by consensus combiners of sequence and structure information. BMC Bioinformatics. 2007;8:201. Epub 2007/06/16. doi: 10.1186/1471-2105-8-201. PubMed PMID: 17570843; PubMed Central PMCID: PMCPMC1913928.

73. Xie S, Li Z, Hu H. Protein secondary structure prediction based on the fuzzy support vector machine with the hyperplane optimization. Gene. 2018;642:74–83. Epub 2017/11/07. doi: 10.1016/j.gene.2017.11.005. PubMed PMID: 29104167.

74. Magnan CN, Baldi P. SSpro/ACCpro 5: almost perfect prediction of protein secondary structure and relative solvent accessibility using profiles, machine learning and structural similarity. Bioinformatics. 2014;30(18):2592–7. Epub 2014/05/27. doi: 10.1093/bioinformatics/btu352. PubMed PMID: 24860169; PubMed Central PMCID: PMCPMC4215083.

75. Heffernan R, Yang Y, Paliwal K, Zhou Y. Capturing non-local interactions by long short-term memory bidirectional recurrent neural networks for improving prediction of protein secondary structure, backbone angles, contact numbers and solvent accessibility. Bioinformatics. 2017;33(18):2842–9. Epub 2017/04/22. doi: 10.1093/bioinformatics/btx218. PubMed PMID: 28430949.

76. Drozdetskiy A, Cole C, Procter J, Barton GJ. JPred4: a protein secondary structure prediction server. Nucleic Acids Res. 2015;43(W1):W389–94. Epub 2015/04/18. doi: 10.1093/nar/gkv332. PubMed PMID: 25883141; PubMed Central PMCID: PMCPMC4489285.

77. Fang C, Shang Y, Xu D. MUFOLD-SS: New deep inception-inside-inception networks for protein secondary structure prediction. Proteins. 2018;86(5):592–8. Epub 2018/03/02. doi: 10.1002/prot.25487. PubMed PMID: 29492997; PubMed Central PMCID: PMCPMC6120586.

78. Torrisi M, Kaleel M, Pollastri G. Porter 5: fast, state-of-the-art ab initio prediction of protein secondary structure in 3 and 8 classes. 2018:289033. doi: 10.1101/289033%J bioRxiv.

79. Wang S, Peng J, Ma J, Xu J. Protein Secondary Structure Prediction Using Deep Convolutional Neural Fields. Sci Rep. 2016;6:18962. Epub 2016/01/12. doi: 10.1038/srep18962. PubMed PMID: 26752681; PubMed Central PMCID: PMCPMC4707437.

80. Zhou J, Wang H, Zhao Z, Xu R, Lu Q. CNNH_PSS: protein 8-class secondary structure prediction by convolutional neural network with highway. BMC Bioinformatics. 2018;19(Suppl 4):60. Epub 2018/05/11. doi: 10.1186/s12859-018-2067-8. PubMed PMID: 29745837; PubMed Central PMCID: PMCPMC5998876.

81. Zhang B, Li J, Lu Q. Prediction of 8-state protein secondary structures by a novel deep learning architecture. BMC Bioinformatics. 2018;19(1):293. Epub 2018/08/05. doi: 10.1186/s12859-018-2280-5. PubMed PMID: 30075707; PubMed Central PMCID: PMCPMC6090794.

82. Heffernan R, Paliwal K, Lyons J, Dehzangi A, Sharma A, Wang J, et al. Improving prediction of secondary structure, local backbone angles, and solvent accessible surface area of proteins by iterative deep learning. Sci Rep. 2015;5:11476. Epub 2015/06/23. doi: 10.1038/srep11476. PubMed PMID: 26098304; PubMed Central PMCID: PMCPMC4476419.

83. Ma Y, Liu Y, Cheng J. Protein Secondary Structure Prediction Based on Data Partition and Semi-Random Subspace Method. Sci Rep. 2018;8(1):9856. Epub 2018/07/01. doi: 10.1038/s41598-018-28084-8. PubMed PMID: 29959372; PubMed Central PMCID: PMCPMC6026213.

84. Hanson J, Paliwal K, Litfin T, Yang Y, Zhou Y. Improving Prediction of Protein Secondary Structure, Backbone Angles, Solvent Accessibility, and Contact Numbers by Using Predicted Contact Maps and an Ensemble of Recurrent and Residual Convolutional Neural Networks. Bioinformatics. 2018. Epub 2018/12/12. doi: 10.1093/bioinformatics/bty1006. PubMed PMID: 30535134.

85. Fourrier L, Benros C, de Brevern AG. Use of a structural alphabet for analysis of short loops connecting repetitive structures. BMC Bioinformatics. 2004;5:58. Epub 2004/05/14. doi: 10.1186/1471-2105-5-58. PubMed PMID: 15140270; PubMed Central PMCID: PMCPMC450294.

86. Ceroni A, Frasconi P, Pollastri G. Learning protein secondary structure from sequential and relational data. Neural Netw. 2005;18(8):1029–39. Epub 2005/09/27. doi: 10.1016/j.neunet.2005.07.001. PubMed PMID: 16182513.

87. Frishman D, Argos P. Knowledge-based protein secondary structure assignment. Proteins. 1995;23(4):566–79. Epub 1995/12/01. doi: 10.1002/prot.340230412. PubMed PMID: 8749853.

88. Yang Y, Heffernan R, Paliwal K, Lyons J, Dehzangi A, Sharma A, et al. SPIDER2: A Package to Predict Secondary Structure, Accessible Surface Area, and Main-Chain Torsional Angles by Deep Neural Networks. Methods Mol Biol. 2017;1484:55–63. Epub 2016/10/28. doi: 10.1007/978-1-4939-6406-2_6. PubMed PMID: 27787820.

89. Altschul SF, Madden TL, Schaffer AA, Zhang J, Zhang Z, Miller W, et al. Gapped BLAST and PSI-BLAST: a new generation of protein database search programs. Nucleic Acids Res. 1997;25(17):3389–402. Epub 1997/09/01. doi: 10.1093/nar/25.17.3389. PubMed PMID: 9254694; PubMed Central PMCID: PMCPMC146917.

90. Rashid S, Saraswathi S, Kloczkowski A, Sundaram S, Kolinski A. Protein secondary structure prediction using a small training set (compact model) combined with a Complex-valued neural network approach. BMC Bioinformatics. 2016;17(1):362. Epub 2016/09/14. doi: 10.1186/s12859-016-1209-0. PubMed PMID: 27618812; PubMed Central PMCID: PMCPMC5020447.

91. Cuff JA, Barton GJ. Evaluation and improvement of multiple sequence methods for protein secondary structure prediction. Proteins. 1999;34(4):508–19. Epub 1999/03/19. PubMed PMID: 10081963.

92. Read RJ, Adams PD, Arendall WB, 3rd, Brunger AT, Emsley P, Joosten RP, et al. A new generation of crystallographic validation tools for the protein data bank. Structure. 2011;19(10):1395–412. Epub 2011/10/18. doi: 10.1016/j.str.2011.08.006. PubMed PMID: 22000512; PubMed Central PMCID: PMCPMC3195755.

93. Remmert M, Biegert A, Hauser A, Soding J. HHblits: lightning-fast iterative protein sequence searching by HMM-HMM alignment. Nat Methods. 2011;9(2):173–5. Epub 2011/12/27. doi: 10.1038/nmeth.1818. PubMed PMID: 22198341.

94. Sievers F, Wilm A, Dineen D, Gibson TJ, Karplus K, Li W, et al. Fast, scalable generation of high-quality protein multiple sequence alignments using Clustal Omega. Mol Syst Biol. 2011;7:539. Epub 2011/10/13. doi: 10.1038/msb.2011.75. PubMed PMID: 21988835; PubMed Central PMCID: PMCPMC3261699.

95. Rost B, Sander C. Prediction of protein secondary structure at better than 70% accuracy. J Mol Biol. 1993;232(2):584–99. Epub 1993/07/20. doi: 10.1006/jmbi.1993.1413. PubMed PMID: 8345525.

96. Frishman D, Argos P. Incorporation of non-local interactions in protein secondary structure prediction from the amino acid sequence. Protein Eng. 1996;9(2):133–42. Epub 1996/02/01. doi: 10.1093/protein/9.2.133. PubMed PMID: 9005434.

97. Cole C, Barber JD, Barton GJ. The Jpred 3 secondary structure prediction server. Nucleic Acids Res. 2008;36(Web Server issue):W197-201. Epub 2008/05/09. doi: 10.1093/nar/gkn238. PubMed PMID: 18463136; PubMed Central PMCID: PMCPMC2447793.

98. Holley LH, Karplus M. Protein secondary structure prediction with a neural network. Proc Natl Acad Sci U S A. 1989;86(1):152–6. Epub 1989/01/01. doi: 10.1073/pnas.86.1.152. PubMed PMID: 2911565; PubMed Central PMCID: PMCPMC286422.

99. Qian N, Sejnowski TJ. Predicting the secondary structure of globular proteins using neural network models. J Mol Biol. 1988;202(4):865–84. Epub 1988/08/20. doi: 10.1016/0022-2836(88)90564-5. PubMed PMID: 3172241.

100. Zhou J, Troyanskaya OG. Deep Supervised and Convolutional Generative Stochastic Network for Protein Secondary Structure Prediction. arXiv e-prints2014.

101. Ting D, Wang G, Shapovalov M, Mitra R, Jordan MI, Dunbrack RL, Jr. Neighbor-dependent Ramachandran probability distributions of amino acids developed from a hierarchical Dirichlet process model. PLoS Comput Biol. 2010;6(4):e1000763. PubMed PMID: 20442867.

102. Shapovalov M, Vucetic S, Dunbrack RL, Jr. A new clustering and nomenclature for beta turns derived from high-resolution protein structures. PLoS Comput Biol. 2019;15(3):e1006844. Epub 2019/03/08. doi: 10.1371/journal.pcbi.1006844. PubMed PMID: 30845191; PubMed Central PMCID: PMCPMC6424458.

103. Li Z, Yu Y. Protein Secondary Structure Prediction Using Cascaded Convolutional and Recurrent Neural Networks. arXiv e-prints2016.

104. Prechelt L. Early Stopping - but when? Neural Networks: Tricks of the Trade, volume 1524 of LNCS, chapter 2: Springer-Verlag; 1997. p. 55-69.

